# Virtual deep brain stimulation: Multiscale co-simulation of a spiking basal ganglia model and a whole-brain mean-field model with The Virtual Brain

**DOI:** 10.1101/2021.05.05.442704

**Authors:** Jil M. Meier, Dionysios Perdikis, André Blickensdörfer, Leon Stefanovski, Qin Liu, Oliver Maith, Helge Ü. Dinkelbach, Javier Baladron, Fred H. Hamker, Petra Ritter

**Author notes:** corresponding authors and; postal address: Charité - Universitätsmedizin Berlin, Charité Campus Mitte (CCM), Brain Simulation Section, Robert-Koch-Platz 4, 10115 Berlin, Germany.

## Abstract

Deep brain stimulation (DBS) has been successfully applied in various neurodegenerative diseases as an effective symptomatic treatment. However, its mechanisms of action within the brain network are still poorly understood. Many virtual DBS models analyze a subnetwork around the basal ganglia and its dynamics as a spiking network with their details validated by experimental data. However, connectomic evidence shows widespread effects of DBS affecting many different cortical and subcortical areas. From a clinical perspective, various effects of DBS besides the motoric impact have been demonstrated. The neuroinformatics platform The Virtual Brain (TVB) offers a modeling framework allowing us to virtually perform stimulation, including DBS, and forecast the outcome from a dynamic systems perspective prior to invasive surgery with DBS lead placement. For an accurate prediction of the effects of DBS, we implement a detailed spiking model of the basal ganglia, which we combine with TVB via our previously developed co-simulation environment. This multiscale co-simulation approach builds on the extensive previous literature of spiking models of the basal ganglia while simultaneously offering a whole-brain perspective on widespread effects of the stimulation going beyond the motor circuit. In the first demonstration of our model, we show that virtual DBS can move the firing rates of a Parkinson’s disease patient’s thalamus - basal ganglia network towards the healthy regime while, at the same time, altering the activity in distributed cortical regions with a pronounced effect in frontal regions. Thus, we provide proof of concept for virtual DBS in a co-simulation environment with TVB. The developed modeling approach has the potential to optimize DBS lead placement and configuration and forecast the success of DBS treatment for individual patients.

**Highlights:**

- We implement and validate a co-simulation approach of a spiking network model for subcortical regions in and around the basal ganglia and interface it with mean-field network models for each cortical region.
- Our simulations are based on a normative connectome including detailed tracts between the cortex and the basal ganglia regions combined with subject-specific optimized weights for a healthy control and a patient with Parkinson’s disease.
- We provide proof of concept by demonstrating that the implemented model shows biologically plausible dynamics during resting state including decreased thalamic activity in the virtual patient and during virtual deep brain stimulation including normalized thalamic activity and distributed altered cortical activity predominantly in frontal regions.
- The presented co-simulation model can be used to tailor deep brain stimulation for individual patients.

## Introduction

Deep brain stimulation (DBS) is a neuromodulation technique that has shown beneficial effects for patients suffering from many different neurological disorders (Horn, 2019; Horn & Fox, 2020). DBS is an essential element in the therapeutic regime for movement disorders like Parkinson’s disease (PD) (Deuschl et al., 2006; Vitek et al., 2020), dystonia (Kupsch et al., 2006) and essential tremor (Koller et al., 1997). It provides a treatment option for selected cases of medication-refractory epilepsy (Salanova et al., 2015) and obsessive-compulsive disorder (OCD) (Anderson & Ahmed, 2003; Franzini et al., 2010; Nuttin et al., 2008). For major depression (Mayberg et al., 2005), Tourette’s syndrome (Ackermans et al., 2011), Huntington’s disease (Gruber et al., 2014) and alcohol addiction (U. J. Müller et al., 2009), DBS has shown first treatment successes and is clinically applied on an experimental basis. Albeit the initial implantation surgery, DBS is a reversible neuromodulation technique, in contrast to a permanent effect after a surgical lesion (Horn & Fox, 2020). Despite the benefits of DBS for many diseases, underlying mechanisms are so far poorly understood. At various scales of the brain, attempts have been made to model the outcome of DBS, from single-neuron to whole-brain models (Humphries et al., 2018). However, a multiscale model to bridge these different scales in a single DBS model has yet to be developed.

The most extensive research for DBS has been performed in movement disorders, which share pathology of the interactions between basal ganglia (BG), thalamus and cortex (Plotkin & Goldberg, 2019). The BG are anatomically defined by the striatum and the pallidum, which can be further separated in globus pallidus internus (GPi) and externus (GPe). Functionally, the regions of the subthalamic nucleus (STN) and the substantia nigra, whose degeneration is a key factor in the pathogenesis of PD (Damier et al., 1999; Fearnley & Lees, 1991), are often included in the BG because of their strong interactions with it (Albin et al., 1989). In the following, the term BG refers to “basal ganglia and related nuclei” (Lanciego et al., 2012) according to the widely used understanding as a functional unit of the extrapyramidal system (Heimer, 1983).

The hypothesis that PD patients often suffer from a decreased activity level in the thalamic region causing the motor function to be impaired, resulting in bradykinesia or akinesia, has a long history (DeLong, 1990; Humphries et al., 2018; Jahanshahi et al., 2015). This decreased activity in the thalamus is probably caused by pathological hyperactivity of the globus pallidus as a failure symptom of the dopaminergic system (Dostrovsky et al., 2002), a theory first formulated by the classical rate model of the BG (Albin et al., 1989). The clinically most relevant stimulation targets for PD are the GPi and STN (Horn & Fox, 2020). It is a common approach to model the neurons of these key regions for DBS as a network, employing mathematical descriptions of neuronal behavior and interactions (Yu et al., 2020). Previous studies found alterations in several important pathways through the BG (direct, indirect and hyperdirect pathway, Figure 1) (Nambu et al., 2002) when simulating DBS. Models of a single neuron and its axon cables propose that STN-DBS causes GPi to fire at a regular frequency (Rubin et al., 2012). An extensive amount of previous literature exists modeling the connection from STN to GPe, the striatal microcircuit and different subparts of the cortico-basal-ganglia-thalamo-cortical loop as spiking networks (Yu et al., 2020). These subnetwork studies often include validation of the model details (parameters as well as results) with experimental data and suggest that STN-DBS changes the efferences of the BG to the thalamus by suppressing the burst firing of the GPi (Guo et al., 2008; Rubin & Terman, 2004). A recent application of subnetwork models shows that STN stimulation causes short-term depression of its own activity (Rosenbaum et al., 2014). This short-term depression theory was validated with empirical data from rodents and primates, where STN-DBS eliminated beta-band oscillations in the GPi in Parkinsonian primate brains (Moran et al., 2012). Another class of models including electrical fields and volume of tissue information proposes that STN stimulation causes heterogeneous effects for different neurons depending on their distance to the electrode (Hahn & McIntyre, 2010; Humphries & Gurney, 2012; McIntyre & Hahn, 2010). These volumetric models explain the heterogeneous effects on the firing rates of GPi neurons, observed experimentally in primates (Hahn et al., 2008; Hashimoto et al., 2003).

**Figure 1:**
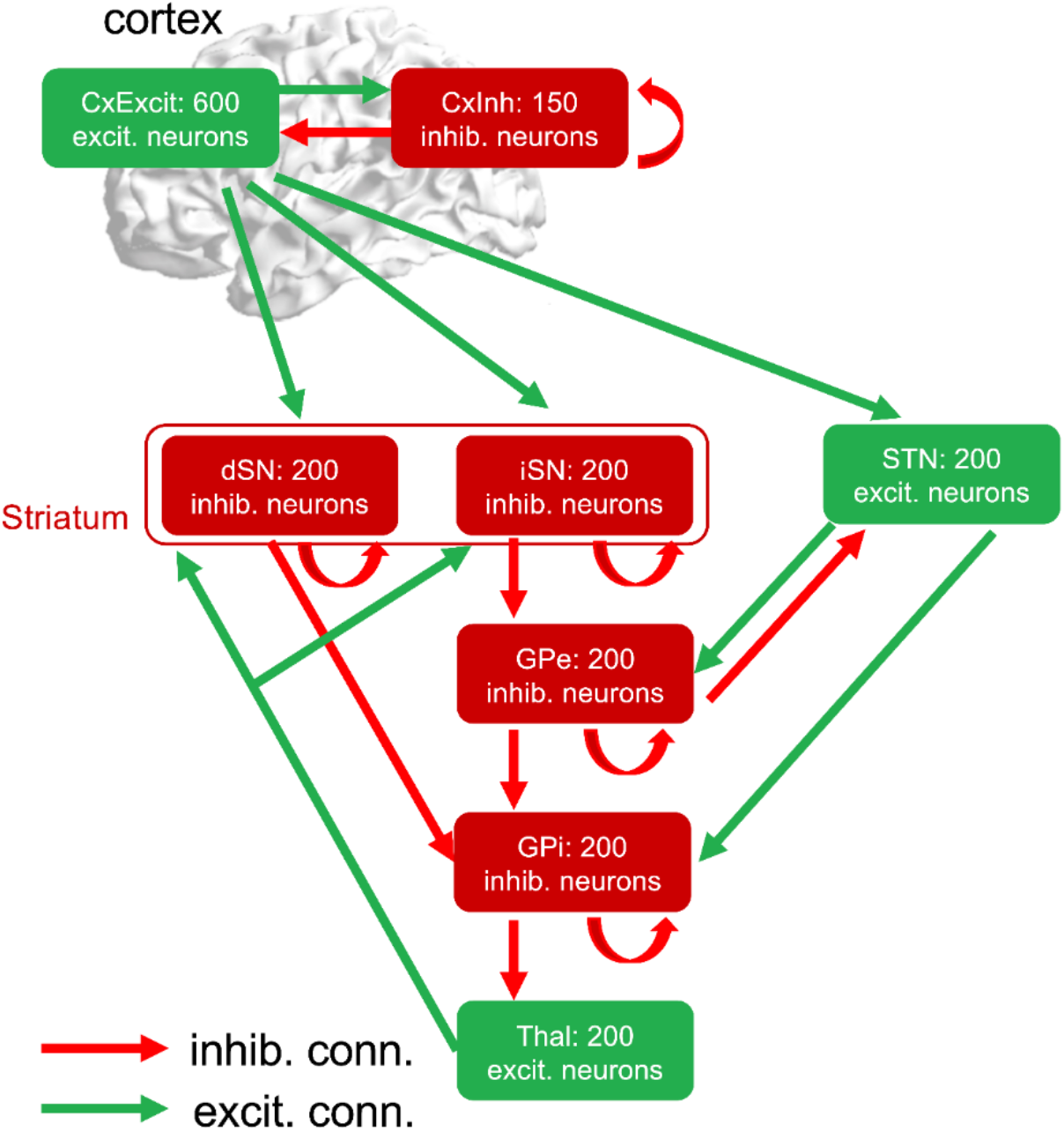
Structure of the basal ganglia spiking model. Previously published detailed basal ganglia (BG) model by (Maith et al., 2020). We implemented this model inside our TVB-ANNarchy framework with the underlying previously optimized connection weights and probabilities for the data of one control and one PD patient (taken from (Maith et al., 2020)). The direct pathway is shown here as the path from the excitatory cortical neurons over the direct striatal projection neurons to the GPi. Similarly, the indirect pathway goes from the cortex, over the indirect spinal projection neurons and the GPe towards the GPi. The third pathway through the BG is the cortex-STN-GPi pathway, which is also called the hyperdirect pathway. CxExcit: excitatory population of the cortex; CxInh: inhibitory population of the cortex; GPi: internal globus pallidus; GPe: external globus pallidus; STN: subthalamic nucleus; dSN: striatum, direct striatal spiny projection neurons; iSN: striatum, indirect striatal spiny projection neurons; Thal: thalamus; excit.: excitatory; inhib.: inhibitory; conn.: connection.

Most previously established models are based on *a priori* assumptions about dynamic changes in PD, i.e., assuming differences between PD and healthy subjects with regard to their functional connectivity strengths or their activity levels of striatal projection neurons (Humphries et al., 2018). Though these assumptions are well justified by empirical findings, they critically influence the model outcomes. In contrast, Hamker and colleagues proposed a data-driven spiking model of the BG (Baladron et al., 2019; Maith et al., 2020), that is a generic BG model has been fit to the individual subject data by optimizing its parameters such that features of the simulated activity correlated with the same features of the measurements. Recently, Maith et al. (2020) fitted this BG model for *20* PD patients after DBS implantation and *15* healthy controls with individual resting-state functional magnetic resonance imaging (fMRI) data. However, the whole cortex was so far modeled as a single spiking network node, lacking a whole-brain perspective. This computational model was implemented with the software Artificial Neural Network architect (ANNarchy), used for spike and rate coding of neuronal populations, as well as a combination of both in a single network (Vitay et al., 2015). Network models in ANNarchy are defined through equations written in “natural language”. ANNarchy has been used to implement models of the BG pathways (Baladron et al., 2019; Baladron & Hamker, 2020; Gönner et al., 2020; Maith et al., 2020; Villagrasa et al., 2018), spatial attention and vision (Bergelt & Hamker, 2019; Jamalian et al., 2017; Larisch et al., 2020) and learning and memory (Gönner et al., 2017; J. Müller et al., 2018; Schmid et al., 2019).

The single-neuron and subnetwork models of the BG successfully suggest underlying mechanisms for the improvement of PD hypokinesia symptoms during DBS. However, they are not sufficient in describing the multitude of other effects that DBS potentially has on PD patients, e.g., rigidity, tremor and cognitive or behavioral changes (Irmen et al., 2019). Therefore, extending local DBS effects of the cortex-BG-thalamus loop towards a large-scale network should be the next goal in understanding DBS effects.

Previous studies explored mean-field approaches simulating the whole-brain perspective for virtual DBS (Saenger et al., 2017; van Hartevelt et al., 2014). Mean-field models make use of a physical simplification to enable simulating the average or so-called mean-field behavior of large populations. Simulating the whole brain with mean-field modeling has shown that DBS brought the patients’ dynamical regime closer to a healthy one (Saenger et al., 2017; van Hartevelt et al., 2014). Specifically, van Hartevelt and colleagues (2014) showed that STN-DBS has widespread structural and functional effects after long-term use analyzing diffusion tensor imaging (DTI) data of a single PD patient. However, they needed to exclude the STN from their analysis as controls were missing MRI data of this region. Saenger et al. (2017) analyzed the fMRI data of 10 PD patients under both conditions DBS switched on and off and performed virtual DBS for different candidate regions, demonstrating their effects for the whole-brain dynamics. This first application of testing DBS effects on the whole-brain dynamics was performed on the group level, while an extension towards the individual level is required before clinical application.

With respect to whole-brain mean-field simulations, The Virtual Brain (TVB, *thevirtualbrain*.*org*) (Ritter et al., 2013; Sanz Leon et al., 2013) offers a neuroinformatics platform to simulate the effects of a virtual DBS. This *in silico* computation of the whole-brain effects of DBS requires only the MRI data of an individual patient as an input. Simulated brain activity with TVB reproduces empirical phenomena accurately over different modalities (Schirner et al., 2018). Applying TVB in combination with simulated stimulation has shown the connection between different stimulation targets and functional resting-state networks based on normative surface-based human brain data (Spiegler et al., 2016). Spiegler and colleagues recently reproduced this finding of activating functional resting-state networks through focal stimulation for the mouse brain, where the results were in line with experimental data of optogenetic stimulation (Spiegler et al., 2020). Transcranial direct current stimulation simulated with TVB based on a normative connectome (Kunze et al., 2016) resembled empirical electroencephalography (EEG) findings. However, virtual DBS has not yet been investigated with TVB.

The different computational studies demonstrating the effects of PD and/or DBS on the BG network, from single-neuron studies to whole-brain networks, exemplify the multiscale nature of this research field (Humphries et al., 2018). So far, the whole-brain DBS modeling literature stands isolated from the extensive literature on spiking neural networks of the BG. Only region-wise properties have been compared. None of the dynamical insights from the spiking network literature have been incorporated into the mean-field modeling approaches of DBS. Therefore, in this study, we aim to demonstrate the framework for a multiscale co-simulation approach of virtual DBS. Our goal is to bridge the microscale of single neurons towards the recorded whole-brain signals in one simulation framework, which permits a holistic and comprehensive integration of existing findings. To run whole-brain mean-field simulations and additionally simulate any region’s fine-scale neuronal dynamics, including spikes generated by inhibitory and excitatory neurons inside the region, we can use the recently developed TVB-multiscale co-simulation toolbox (Schirner et al., 2021). TVB-multiscale extends TVB to perform multiscale co-simulations, whereby most of the nodes are simulated with TVB as mean-field models, and a few selected nodes are modeled as spiking networks by another suitable simulator.

In this study, we combine the detailed spiking network model by Maith et al. (2020) for the BG with mean-field simulations in TVB for all cortical regions. We interface the spiking network software ANNarchy with TVB to build the TVB-ANNarchy co-simulation framework (Schirner et al., 2021). As an underlying connection between BG and cortical regions, we utilize a recently published normative connectivity atlas of these tracts (Petersen et al., 2019) and combine it with individually - that is subject-specific - fitted probabilities and weights from Maith et al. (2020) for the connections among the BG regions. As a first proof of concept, we simulate resting-state conditions for an exemplary control and PD patient network and perform virtual DBS targeting STN and GPi in the patient network. Next, we validate our model by comparing the effects of virtual DBS against results from literature. Our study addresses the following limitations of previous whole-brain DBS modeling studies:

1. We incorporate a previously validated spiking network model of the subnetwork of the BG within our whole-brain modeling.
2. We use an underlying (normative) connectome, which includes the STN, and combine it with individually fitted connectivity data to create an individual patient and control multiscale network.

In this way, we offer a computational model that holds the potential to be easily translated towards the individual patient level and used as a ‘sandbox’ model before future DBS surgeries.

## Materials and Methods

### Spiking network model for the basal ganglia

The spiking network model and its dynamics (including parameters) were taken from a previous publication (Maith et al., 2020) (Figure 1). Eight neuronal populations were included, each with different properties. The cortex consisted of 600 excitatory neurons coupled with 150 inhibitory neurons (possessing a self-inhibitory connection). From the excitatory population of the cortex, spikes were transmitted to the STN, as well as the striatum. The striatum was modeled with two different inhibitory neuronal populations, the direct (dSN) and the indirect (iSN) striatal spiny projection neurons, each with a self-inhibitory connection. The GPe and GPi were each represented by inhibitory neurons with a self-inhibitory connection. The thalamus was also modeled as a spiking network node.

Each spiking network population was modeled by an enhanced version of the Izhikevich model (Izhikevich, 2004; Maith et al., 2020). For details of this previously published model, we refer to the Supplementary Material. Maith et al. (2020) optimized the connection probabilities and weights between the nodes to fit empirical fMRI blood-oxygen-level-dependent (BOLD) signal correlation data for each individual and each hemisphere separately. We used this optimized data from one of the controls and one of the patients (left hemisphere only). We selected these subjects as representatives of their groups because their regional firing rates were close to the respective mean values.

### Multiscale co-simulation of TVB and ANNarchy

Every node in the TVB network represents a brain region and its dynamics are simulated with a mean-field approximation. The nodes are connected with weights and delays (computed from tract lengths given a transmission speed) that can be determined for individual subjects employing DTI. As a mean-field model for the cortical regions, we chose the reduced Wong-Wang-model (Deco et al., 2013), which is often used to replicate fMRI data (Aerts et al., 2018; Klein et al., 2021) (details in the Supplementary Material). For an overview of all variables used in this study, we refer to Supplementary Table 1. In the TVB-multiscale framework (Schirner et al., 2021), co-simulation is based on the concept of TVB “proxy” nodes that are created inside the spiking network (Figure 2). TVB “proxy” nodes are either stimulating devices, thereby mimicking TVB cortex node dynamics (i.e., mean-field spiking rates) and coupling to the spiking nodes, or output (e.g., recording) devices, thereby extracting spiking dynamics to be transmitted to TVB. Thus, TVB and the spiking network simulator communicate on the level of neuronal populations’ mean-field activities. TVB-multiscale, which is continuously expanding, is freely available on github (github.com/the-virtual-brain/tvb-multiscale) and interfaces TVB with different spiking network simulators (currently Neural Simulation Technology (NEST) (Eppler et al., 2008) and ANNarchy).

**Figure 2:**
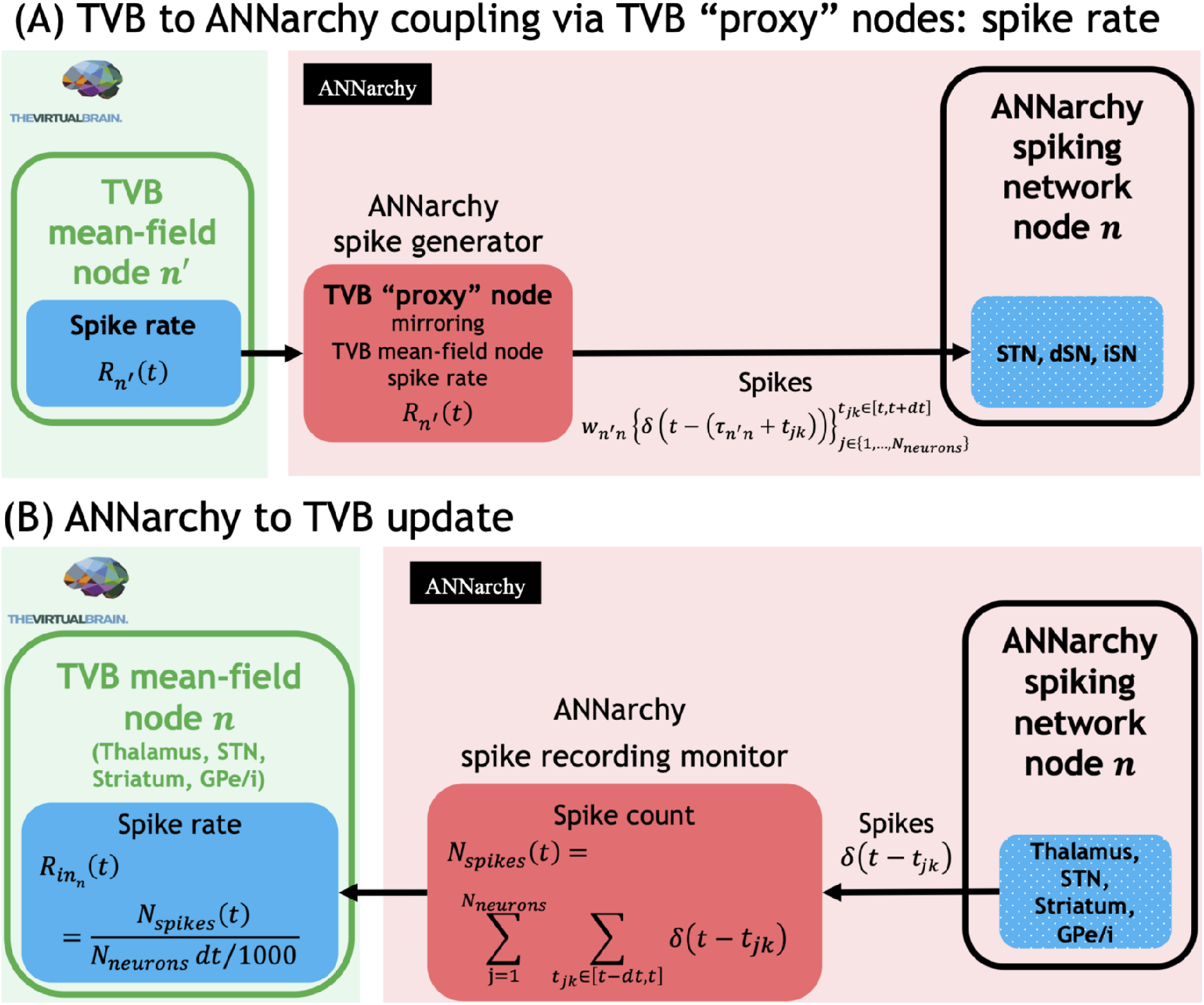
Implementation of the interface for the multiscale model. **(A)** TVB to ANNarchy coupling is channeled via TVB “proxy” nodes in the ANNarchy network, implemented as neuronal populations generating correlated spike trains. Thus, the instantaneous mean-field spike rate is transformed into individual neurons’ spike trains for the respective time interval. **(B)** ANNarchy to TVB state update via the ANNarchy monitors, which record the spikes for each time interval [t −dt,t]to compute the population’s spike rate. This spike rate then overwrites the respective TVB state variable.

Since the previous BG model implementation was fitted with empirical data using ANNarchy (Maith et al., 2020), we built an interface between ANNarchy and TVB. We developed python code to incorporate the ANNarchy simulator into TVB-multiscale (details in the Supplementary Material). We validated our implementation of the spiking network by Maith et al. (2020) against the authors’ original ANNarchy code by performing short simulations without noise for the two selected subjects (Supplementary Table 3).

Each TVB cortex mean-field node *n*′ (prime notation for nodes modeled only as mean-fields nodes in TVB) couples to a node *n* modeled in ANNarchy (notation without prime for the spiking regions) via the instantaneous spike rate variable *R* _*n*′_(*t*), which drives a population of *N*_*neurons*_=600 neurons (same size as for the excitatory cortex node of the spiking network by Maith et al. (2020)) generating correlated spike trains

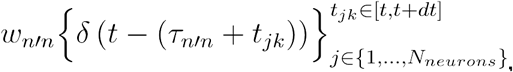

where *t*_*jk*_ stands for the spike time *t*_*k*_of the neuron with index *j* in the population of the “proxy” node *n*′ and *δ* is the Kronecker delta. The generated spikes were weighted by *w* _*n′n*_ and delayed by *τ* _*n′n*_ based on the TVB connectome and the optimized weights for each subject (see below). For details of the spike trains’ generation, we refer to the Supplementary Material.

In the other direction, each node *n* modeled in ANNarchy updates the state of the corresponding TVB mean-field node *n* since it is still represented in the TVB model and couples to TVB nodes *n*′. The update utilizes an ANNarchy monitor that records spikes for each TVB time step. The recorded spikes are converted to an instantaneous population mean rate that overwrites an auxiliary TVB state variable, called the input rate 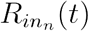. The latter drives a linear integration equation of another auxiliary TVB state variable, named integrated rate 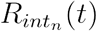, which, in its turn, acts as a smoothing low pass filter

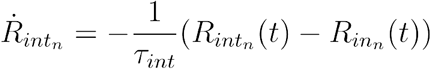

to have time series similar to the TVB mean-field ones, where *τ*_*int*_ = 100*ms* is the time constant of the integration and

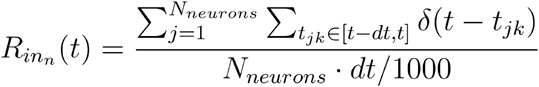

provides the number of spikes per second. Finally, the integrated rate 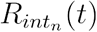 overwrites the state variable *R*_*n*_(*t*) of the TVB model. All the rest of the TVB mean-field nodes *n*′ follow the equations of the mean-field model described in the Supplementary Material. We simulated two ANNarchy time steps (of .0.05*ms*) for every TVB time step (*dt*=0.1*ms*).

### Underlying connectivity

To connect the TVB nodes and the spiking network simulator, we needed to assign connectivity weights for the paths between the BG regions and the cortex. Acquiring accurate data for those tracts is challenging because structural MRI data inherits many limitations (Jones et al., 2013; Thomas et al., 2014). Recently, Petersen et al. (2019) published a state-of-the-art axonal pathway atlas for the human brain that combines previous results from histological and imaging data literature with expert knowledge of neuroanatomists and brain-imaging scientists who collaborated on defining those tracts applying a holographic visualization technique (Petersen et al., 2019) (details to be found in the Supplementary Material). We used this normative tract data by Petersen et al. (2019) to include a fine-grained parcellation for the BG and the thalamus (based on CIT-168 brain atlas (Pauli et al., 2018)) and detailed data of their pathways to and from the cortical regions because of its current use for clinical DBS planning (Noecker et al., 2021). Whereas Maith et al. (2020) used the motoric parts of the BG regions only (Horn et al., 2019), we used the complete BG regions as a first approach. For the cortex, the automated anatomical labeling (AAL) atlas parcellation was applied (Rolls et al., 2015; Tzourio-Mazoyer et al., 2002). The files were transformed to the DBS Intrinsic Template Atlas (DISTAL) space (Ewert et al., 2018) and the number of streamlines between each region pair was counted. This procedure resulted in a whole-brain matrix for the pathways between the cortex and the BG structures.

Some additional preparation steps have been performed on the connectome. As a first demonstration and because Maith et al. (2020) also treated the hemispheres in isolation, we focused on the left hemisphere only. Thus, all regions belonging to the right hemisphere and the vermis have been deleted from the connectome together with all their connections. Additionally, the connections from the inhibitory neuronal populations in the BG (GPe, GPi and striatum) to any cortical regions have been set to zero as they are not biologically plausible from a functional perspective of movement regulation, leaving in this direction only the connections from the thalamus and the STN to the cortex. The resulting connectome included 57 regions (for a list of all included regions: Supplementary Table 4). Its weights were normalized by the sum of all the incoming connection weights of the corresponding region and by the 99^th^ percentile of all weights.

The previous work of Maith et al. (2020) optimized the connection probabilities and weights among BG regions per individual to best fit the empirical fMRI data. To personalize the normative connectome, we replaced the network among the BG and thalamus regions with the optimized weights computed by Maith et al. (2020) for the control subject and the PD patient, respectively (Figure 3). This ‘hybrid’ connectome constituted normative connectome weights among the cortex regions and between cortex and BG but included individually fitted connection weights and probabilities for the spiking network of the BG. For the presentation and for determining the couplings between the two scales, we adjusted the normative weights to be in the same range of values as the optimized connection weights by scaling them with the ratio *C*_*norm*_ between the 95^th^ percentiles of both weight distributions. The global coupling *G* of the TVB mean-field model was set for each subject to *G*=15/ *C*_*norm*_, i.e., we are canceling the above normalization for the weights among the TVB nodes (see next section for the exact procedure of selecting the value of 15). The conduction speed was set to 4*m*/*s*, thus, determining the time delay of couplings among all nodes of the multiscale model. The tract lengths among all regions were approximated by the Euclidean distance between their center coordinates (Supplementary Figure 2).

**Figure 3:**
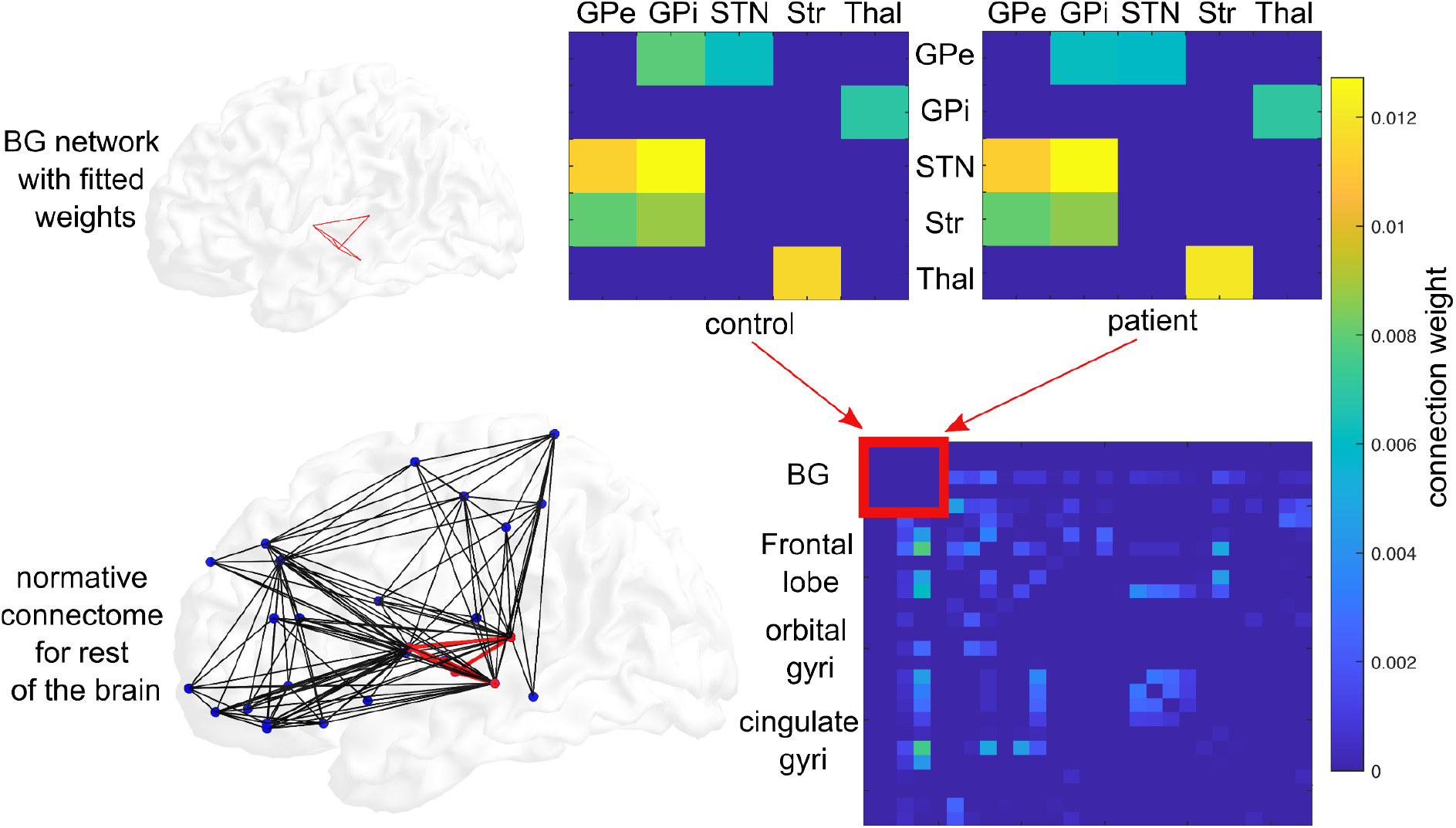
Underlying connectome. The optimally fitted connectivity data from Maith et al. (2020) for the left hemisphere of the analyzed control and patient (upper panels) overrode the within-BG connection weights inside the connectome (red square) based on (Petersen et al., 2019). Each entry in any of the three colored matrices represents the normalized number of streamlines that start in the region marked on the vertical axis and end in the region marked on the horizontal axis. The brain network in the lower panel shows all connections taken from the individually fitted weights and the BG regions in red and the other regions in blue with the connections taken from the normative connectome of the atlas by Petersen et al. (2019) represented in black. For visualization purposes, all isolated nodes have been disregarded. GPi: internal globus pallidus; GPe: external globus pallidus; STN: subthalamic nucleus; dSN: striatum, direct striatal spiny projection neurons; iSN: striatum, indirect striatal spiny projection neurons; Thal: thalamus; Str: striatum.

### Fitting the co-simulation model to individual dynamics

We implemented the previous BG model by Maith et al. (2020) inside our TVB-ANNarchy framework (Figure 4). For the multiscale model (“TVB-cortex model”), we replaced the spiking node “cortex” with the whole brain connectomic model in TVB (Figure 5). However, the input from the multitude of the TVB mean-field nodes leads to different driving dynamics of the spiking network than in Maith et al. (2020). We aimed for TVB driving dynamics that would exhibit (a) a mean firing rate across all TVB nodes of 10 −15*Hz* similar to motor cortex neurons at rest (Velliste et al., 2014), where the range of rate values across TVB regions is determined by the structural connectome; (b) low amplitude random fluctuations of the rate around the equilibrium point of the above mean rate, resembling the rate dynamics of the cortex node in Maith et al. (2020) (c) some correlation among the neurons’ spiking, which in Maith et al. (2020) is due to the internal connectivity of the spiking cortex node populations. We set the operation point of the TVB mean-field network by progressively increasing global coupling *G* until an equilibrium point was reached (for *G*=15/ *C*_*norm*_) with a mean firing rate across the whole TVB brain approaching 15*Hz* from below via a few “trial and error” simulations. After the equilibrium point was approximated, we increased the additive white noise to a standard deviation of *10*^*-4*^ allowing small fluctuations around the equilibrium point without changing the pattern of nodes with higher firing rates (Supplementary Figure 3 displays a characteristic TVB time series during co-simulation).

**Figure 4:**
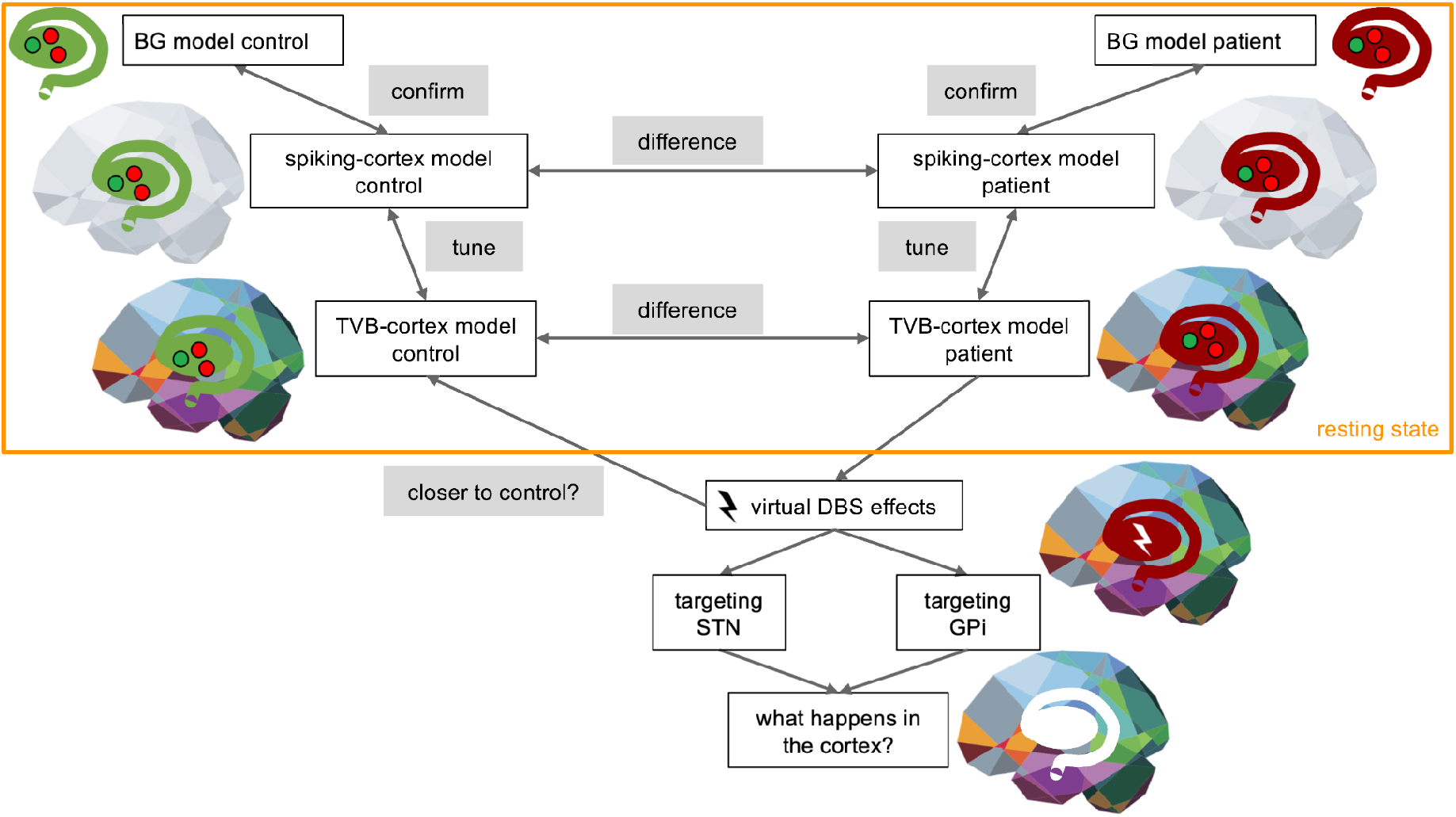
Schematic overview of our study design. The basal ganglia (BG) model of one control and one Parkinson’s disease patient were taken from the previous study (Maith et al., 2020). Next, we implemented the previous model inside our TVB-ANNarchy framework, not yet activating TVB, the so-called “spiking-cortex model”. We confirmed that this implementation reaches similar firing rates as the one from the previous study (step “confirm”). As a second step, we replaced the spiking-cortex node with mean-field simulations using TVB. To stay in the range of the previously confirmed firing rates for the BG regions, we fine-tuned the connection weights from TVB to ANNarchy for the TVB-cortex models of the control and the patient. So far, all of the described modeling steps were taken in resting-state conditions. As a third step, we stimulated the STN and the GPi as two frequently targeted regions virtually (virtual DBS) and analyzed the effects for the BG spiking network as well as for the cortical regions. We analyzed whether virtual DBS could bring the patient’s brain dynamics closer to the healthy one. Whenever there is a brain next to the model (even when it is grayed out), the simulation took place inside the TVB-ANNarchy environment.

**Figure 5:**
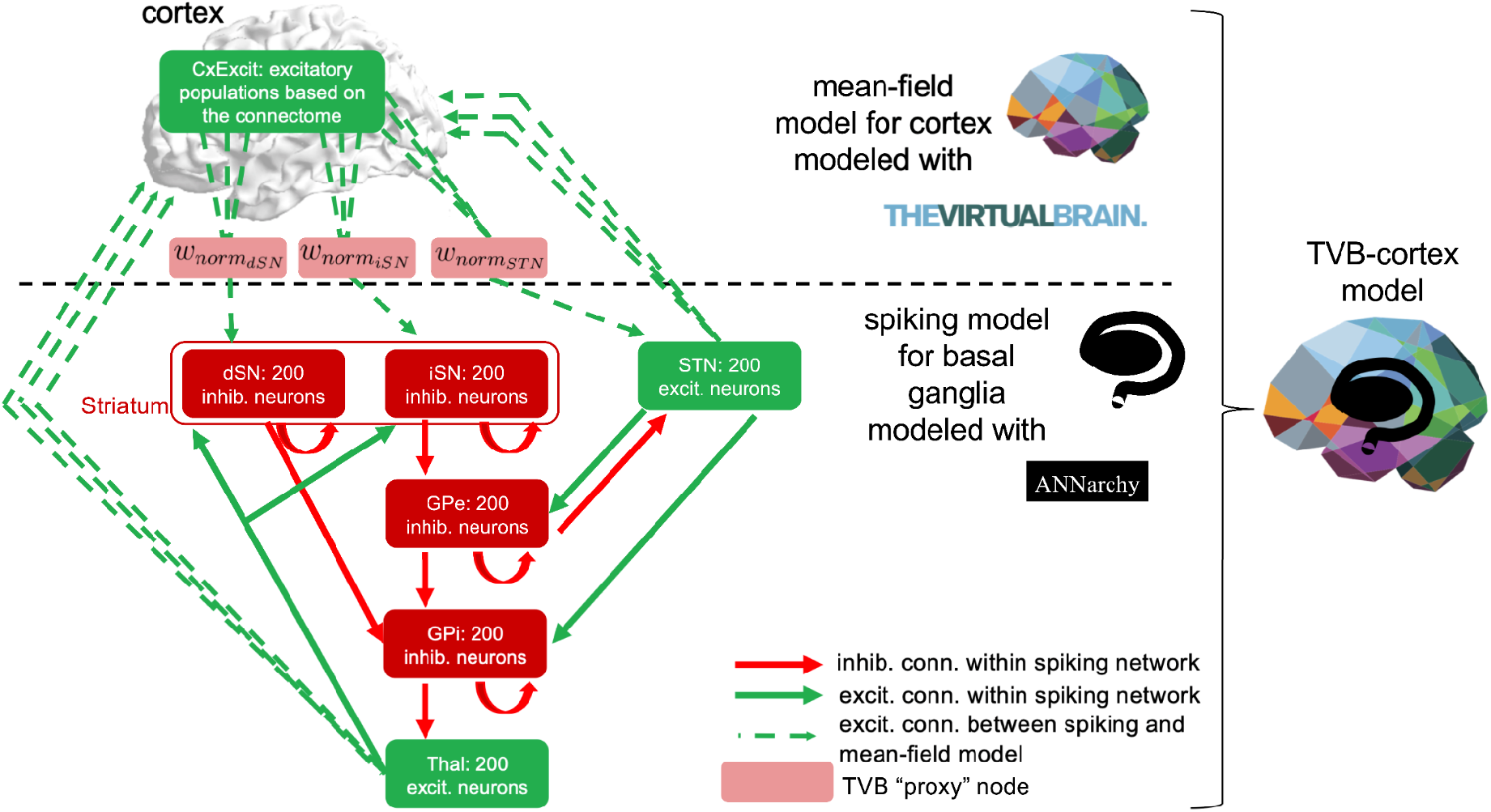
Structure of the co-simulation model. The cortex node was replaced by a whole-brain network simulated with The Virtual Brain. The interactions among the cortical regions were simulated with a mean-field model and The Virtual Brain (TVB). The computational spiking model was simulated with ANNarchy. Together, these two models for the multiscale network, the so-called TVB-cortex model. Interactions between the mean-field and the spiking model were defined by the connection weights of the underlying connectome between all involved region pairs. Connections from cortical regions toward the spiking network (i.e., in our case toward dSN, iSN and STN) were bundled together for each of the regions receiving input from the cortex. In addition, the bundled connections were weighted with the interface weights 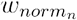, n ∈{dSN, iSN, STN} to regulate the incoming driving activity of the network. CxExcit: excitatory populations of the cortex; GPi: internal globus pallidus; GPe: external globus pallidus; STN: subthalamic nucleus; dSN: striatum, direct striatal spiny projection neurons; iSN: striatum, indirect striatal spiny projection neurons; Thal: thalamus; excit.: excitatory; inhib.: inhibitory; conn.: connection.

For the multiscale TVB-cortex model, the three connections from cortex to STN, dSN and iSN were substituted by the respective set of connections from each of the corresponding TVB nodes (Figure 5). For scaling these connections, we created a “spiking-cortex model” by substituting the cortex node of the network from Maith et al. (2020) with an ANNarchy spike generator identical to the one used as TVB “proxy” nodes (Supplementary Material). The spiking-cortex model acted as the “bridge” between the noisy TVB cortex driving the multiscale model and the Izhikevich population spiking cortex of Maith et al. (2020). With this model, we performed resting-state simulations for both subjects. Then, we tuned - again via a few “trial and error” co-simulations - three interface factors 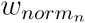,*n* ∈{*iSN,dSN,STN*}, scaling all TVB connections to STN, dSN and iSN, respectively, to approximate the mean population rates of the spiking-cortex model (Figure 5, Supplementary Table 1). These interface factors multiply the TVB connectome weights *C* _*n*′*n*_ resulting in the interface weights 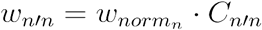 (Figure 2A). This step was also taken to ensure that dSN and iSN can receive different input while the striatum connectivity is equally strong for them. This way, the sum across all TVB nodes *n*′ is the resulting total weight of the cortex input to the BG spiking populations, which then has an effect close to the one of the optimized weights in Maith et al. (2020). The final mean firing rates were approximated within 1*Hz* for all spiking populations except for the thalamus of the control network, which was within 2*Hz* (Supplementary Table 5, Supplementary Figure 4).

For the results of the TVB-cortex simulations, we simulated each condition *10* times, randomly selecting initial conditions for the TVB state from a normal distribution with mean equal to the initial conditions used originally for fitting the resting-state simulations and standard deviation *0*.*1* (Supplementary Material). The simulation length for all of our simulations was 1500*ms*. After each simulation, we computed the mean firing rate over the last 1000*ms*.

### Implementation of the DBS stimulus

Besides the resting-state co-simulations, we applied a stimulus to our multiscale model (starting at 400*ms* and lasting till the end) inside either GPi or STN as possible target regions (Figure 5). We simulated the propagation of these stimuli and the whole-brain response to them to provide a first proof of concept of the possibilities of this kind of multiscale modeling. We tested the virtual DBS stimuli within the spiking-cortex network and the TVB-cortex simulation model.

To the GPi, we applied a constant inhibitory current stimulus of an amplitude of −10*Pa* aiming at reducing its firing rate and therefore disinhibiting the thalamus. For the other DBS simulations, we applied two realistic stimuli to STN, a monophasic and biphasic pulse-like current because the former is the most commonly implemented stimulus in previous DBS simulation studies (Yu et al., 2020) and the latter is used in clinical practice (Krauss et al., 2021) (Figure 6). The monophasic stimulus is adapted from (Michmizos & Nikita, 2011) and the biphasic stimulus is similar to the one used in (Liu et al., 2020) (details in the Supplementary Material).

**Figure 6:**
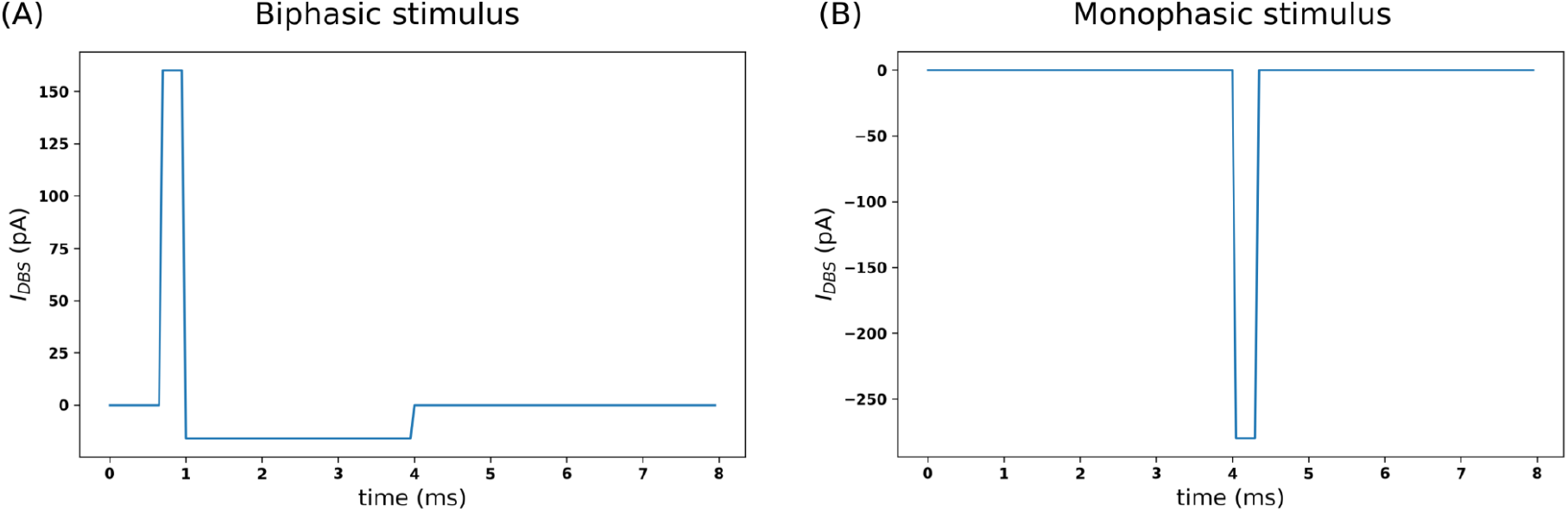
STN-DBS stimulus patterns. One cycle of the **(A)** biphasic and **(B)** monophasic stimuli applied for the virtual DBS targeting the STN region.

### Effects of the stimuli on cortical regions

To investigate the effects of the different stimuli on the cortex, we compared our resting-state TVB-cortex simulations with the simulations including the stimuli for the patient network. We also investigated cortical differences between the resting-state condition of the control and the patient. For these comparisons, we calculated the region-wise difference of cortical firing rates between simulations. Firing rates were averaged over the last 1000*ms* of a simulation. In the resting-state case of comparing the patient and the control, we subtracted the average firing rates of the control’s resting-state simulation from the ones obtained with the patient network. For evaluating the cortical effects of the different stimuli, we subtracted the resting-state average firing rates from the ones of the stimulus-induced time series. In addition, the resulting regional differences were normalized by the mean difference over the obtained regional differences, for each subtraction separately.

All of our code is publicly available (https://github.com/the-virtual-brain/tvb-multiscale/tree/Meier_etal_ExpNeur2021).

## Results

For the whole-brain co-simulation model, we visualized the raster plots of the spiking-network regions for the four conditions, resting-state, GPi-DBS, STN-DBS with a monophasic and STN-DBS with a biphasic stimulus (Figure 7). Comparing the resting-state firing rates, the largest difference between the patient and the control can be found in the thalamus (Figure 8 and Supplementary Table 4). The stimuli caused the biggest changes in firing rate in the stimulated regions themselves (STN or GPi, respectively) and also in the thalamus (Figure 8 and Supplementary Table 4). The GPi-DBS simulation induced disinhibition of the thalamus from the GPi, allowing the thalamus to fire more than in the resting-state condition. Both STN-DBS simulations, however, also showed increased thalamic activity compared to the resting state but together with an increased firing in the GPi (Figure 8). Compared with the resting-state firing of the control, the patient seems to come closer to the rates of the control in multiple regions of the BG during all DBS scenarios (Figure 8). The common mechanism over all three stimulation protocols was the increase in thalamic activity. Thus, the thalamus firing rate seems to “normalize” toward the healthy regime during virtual DBS.

**Figure 7:**
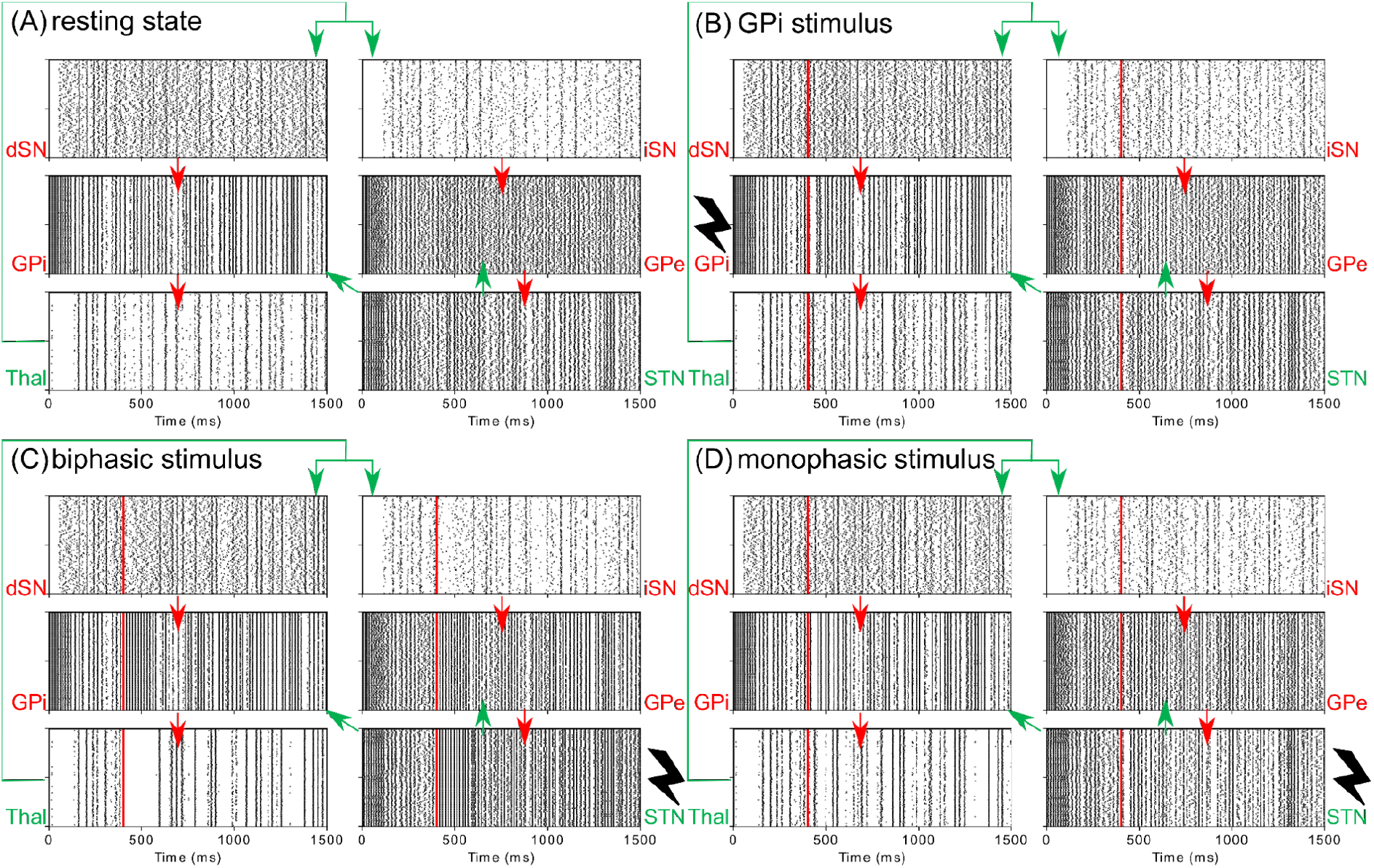
Resting-state and DBS simulation results of the full co-simulation model implemented in TVB-ANNarchy for the patient network. The cortex is represented by the full-scale TVB model with a neural-mass model for each region, i.e., the TVB-cortex model results are displayed. The raster plots of the regions simulated with ANNarchy are shown here. On the y axis, the 200 neurons are listed in the respective region. Each dot in the raster plot represents a spike time of an individual neuron. Vertical black bars in the raster plot, thus, represent synchronous firing activity of all neurons. Mean firing rates are calculated based on the last 1000ms of each simulation. **(A)** Results of the resting-state simulation for the patient’s network. **(B)** Results of a virtual DBS simulation targeting the GPi with an inhibitory continuous current stimulus. The disinhibiting effect of the GPi stimulation (from 400ms onwards) towards the thalamic activity can be observed in the visualized raster plots. **(C)** Results of a virtual DBS simulation targeting the STN with a biphasic stimulus. **(D)** Results of a virtual DBS simulation targeting the STN with a monophasic stimulus. The red vertical lines in the plots represent the start of the respective stimulus. Red (green) arrows visualize inhibitory (excitatory) connections among the regions. Region names written in red (green) color-code an inhibitory (excitatory) population. GPi: internal globus pallidus; GPe: external globus pallidus; STN: subthalamic nucleus; dSN: striatum, direct striatal spiny projection neurons; iSN: striatum, indirect striatal spiny projection neurons; Thal: thalamus.

**Figure 8:**
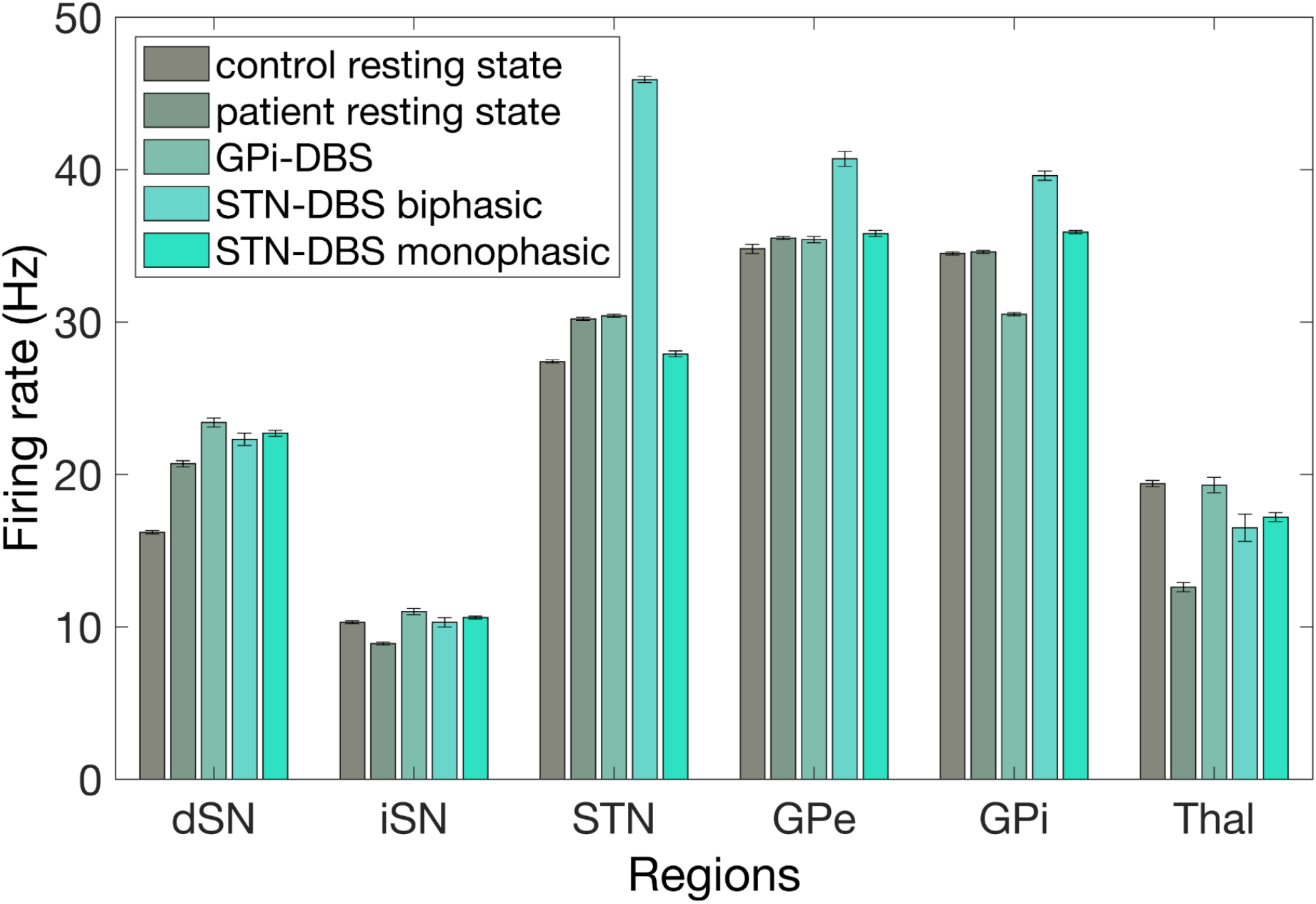
Average firing rates obtained by different simulations of the TVB-cortex model. For each of the six spiking regions, the first and second bar represent the resting-state condition for the control and the patient, respectively. The latter three bars correspond to the three virtual DBS simulations, i.e., GPi-DBS, STN-DBS applying a biphasic and a monophasic stimulus. The height of the bar represents the firing rate (in Hz) averaged over the last 1000ms of the respective simulation and over the 10 simulation repetitions. The error bars have the length of twice the standard deviation over the average firing rates obtained over these 10 repetitions. For the thalamic firing rate, we observe a lower firing rate for the resting-state simulation of the patient compared to the control. After stimulation, the firing rate of the thalamus increased. GPi: internal globus pallidus; GPe: external globus pallidus; STN: subthalamic nucleus; dSN: striatum, direct striatal spiny projection neurons; iSN: striatum, indirect striatal spiny projection neurons; Thal: thalamus.

For the spiking-cortex model, the effects of the stimuli could not be traced further towards the cortical regions since this model is isolated and lacks embedding in the whole-brain network. Comparing the resting-state activities of the cortical regions between patient and control showed an increased average firing rate in the frontal regions and a decreased firing rate in the postcentral gyrus for the patient (Figure 9A). Regarding the cortical effects of stimulation for the TVB-cortex model, we plotted the differences measured by the average firing rate between the resting-state and each stimulus simulation per cortical region on the template brain in Figure 9B-D. In all three virtual DBS simulations, the largest induced changes among the cortical regions were found in the frontal lobe and additionally in the postcentral gyrus for the monophasic STN stimulus. Altered levels of firing rates induced in the GPi and STN by the stimuli appear to be conveyed towards these cortical regions altering their activity with respect to the resting state. Concerning the specific regions, the middle frontal gyrus and insula were for all three stimuli among the top five regions regarding the most increased firing rates induced by the stimulus. Interestingly, only the STN monophasic stimulus created a slight reduction of firing rate in the supplementary motor area.

**Figure 9:**
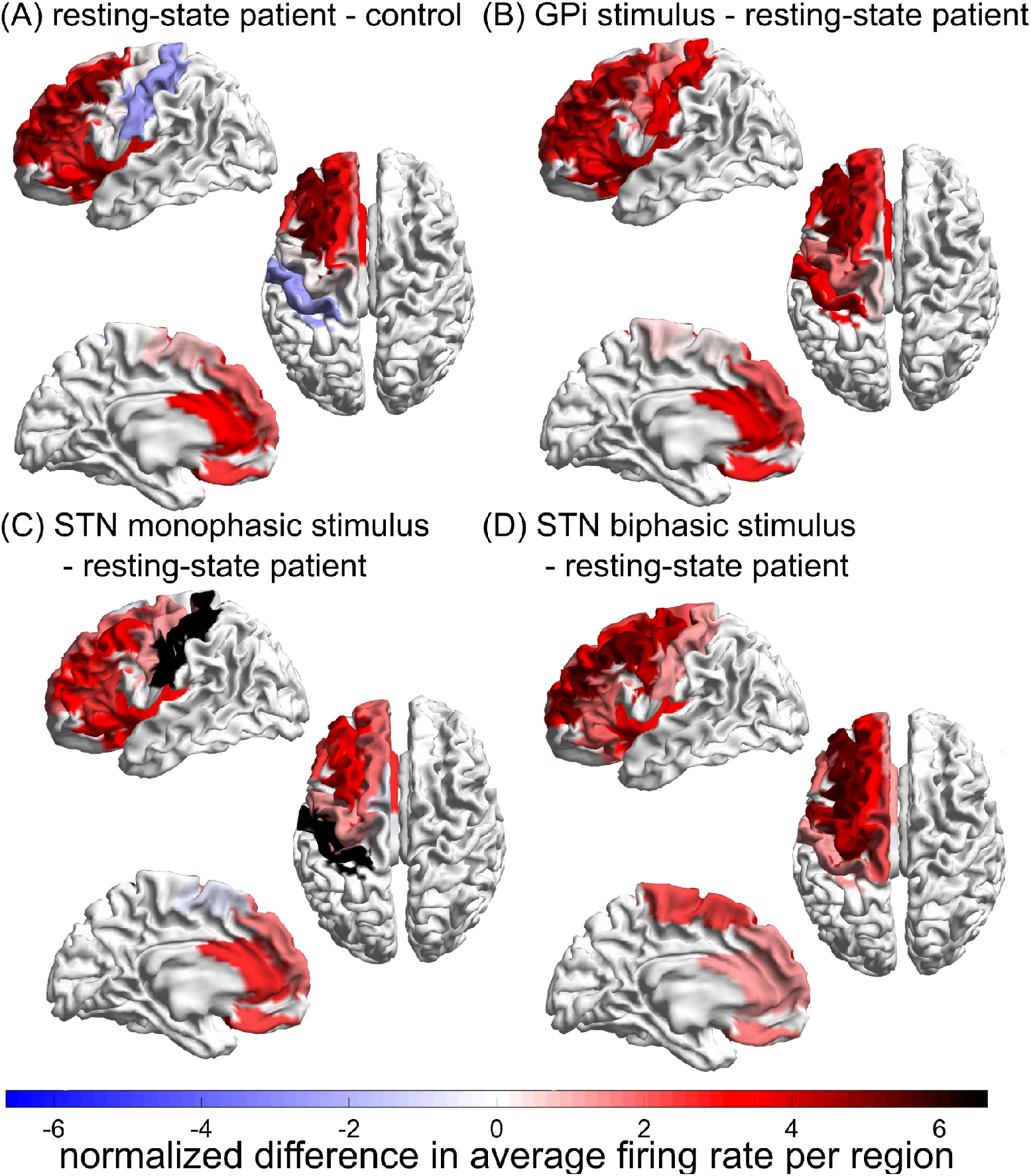
Effects of the stimuli on cortical regions. We plotted the differences in averaged firing rate over the last 1000ms of the simulation time on the template brain, subtracting the average rate obtained from one condition from the other. In addition, the resulting regional differences were normalized by the mean difference over the obtained regional differences for each subtraction separately. **(A)** The normalized difference in average firing rates is shown when subtracting the resting-state simulation results of the control from the ones of the patient. **(B)** The normalized difference in average firing rates is shown when subtracting the GPi stimulus simulation results from the resting-state simulation results of the patient. STN biphasic and **(C)** The normalized difference in average firing rates is shown when subtracting the STN monophasic stimulus simulation results from the resting-state simulation results of the patient. **(D)** The normalized difference in average firing rates is shown when subtracting the STN biphasic stimulus simulation results from the resting-state simulation results of the patient. Thus, red (blue) colors indicate an increased (decreased) average firing rate of that specific region compared with the resting-state condition of the control (A) or the resting-state condition of the patient (B-D). Since our simulations are limited to the left hemisphere, we visualized the differences only for the left hemisphere.

## Discussion

In this study, we introduced a multiscale modeling strategy for the brain network, which allows to model the spiking network dynamics of the BG subnetwork in detail while simultaneously offering a whole-brain perspective of the evolving dynamics. We showed a first proof of concept that this new resulting TVB-multiscale model generates biologically plausible activity in resting state and after virtual DBS. This model has the potential to forecast DBS effects for different locations and different configurations on an individual patient level.

Our presented results show that the DBS stimulus introduced on our patient network causes disinhibition of the thalamus, leading to an increased firing rate during stimulation compared to resting state. Even though our results are in line with the hypotheses formulated by the classical rate model (Albin et al., 1989), conflicting evidence from clinical studies suggests a broader perspective as reduced thalamic activity alone neither explains all symptoms of PD nor all existing therapeutic effects (Eisinger et al., 2019; Marsden & Obeso, 1994; Rodriguez-Oroz et al., 2009). As for the direct effect of the stimulus on the target region, the recent theory of short-term depression states that STN-DBS blocks the transfer of low-frequency oscillations downstream, e.g., towards GPe and GPi, and brings the thalamic activity back to healthy functioning (Humphries et al., 2018). Other theories exist about the effects of the DBS stimulus being of excitatory, inhibitory or disruptive nature on its neighboring areas and a consensus has yet to be reached in this research field (Chiken & Nambu, 2016). Still, our results show that the thalamic activity was brought back to healthy functioning by DBS, which is in line with the general mechanism of DBS (Humphries et al., 2018).

The increased firing rates of the thalamus during our STN-DBS simulations are not caused by decreased GPi activity, which cannot be explained by the classical direct/indirect pathway model of BG. Empirical evidence supports the observed increased firing rates of GPi during STN stimulation (Reese et al., 2011), which were assumed to overwrite pathological activity patterns. One recent computational modeling study with optogenetic data of rodents has shown that increased GPi activity, when synchronized, is able to drive excitatory thalamic responses despite the inhibitory nature of the connection (Liu et al., unpublished results). The proposed underlying mechanism is that bursts of inhibition from GPi to thalamus can cause hyperpolarization and then post-inhibitory rebound firings of thalamus neurons. Post-inhibition spikes or bursts are characteristic behavior of the Izhikevich neuronal model used in this study (Izhikevich, 2004). Taking this unclear mechanism of pacing into account, our model provides computational evidence supporting a more connectomic effect leading to thalamic activation.

There have not been previous studies of multiscale co-simulation of DBS. PD is a multiscale disease (Kerr et al., 2013) with pathological mechanisms at many different scales, from deterioration observed in single neurons up to large-scale brain dynamics. Thus, in the attempt of modeling the broad perspective of potential treatment effects, one should also no longer focus on a single scale. One previous study embedded a spiking network for BG regions inside a neural field model for the cortex (Kerr et al., 2013). However, this previous modeling strategy did not subdivide the cortex mean-field model further into separate regions nor did the authors simulate DBS.

Compared to spiking models that encompass the BG regions only, our presented model can show whole-brain effects of stimulation going beyond the motor cortex. The presented results show an increase in overall activity in cortical regions for all of the three applied stimuli. This result is in line with the theory that PD patients have lower thalamic activity and, thus, a weaker driving activity from the thalamus towards the cortex. Subsequently, the cortex reacts with an increase of activity to the DBS-induced disinhibition of the thalamus. The frontal regions and the insula seem to be most impacted by all three different stimuli, measured by an increase in firing rate. The insula is linked strongly with non-motor symptoms in PD (Christopher et al., 2014) and a previous study reported a BOLD signal increase in the insula during STN-DBS (Kahan et al., 2012). The middle and inferior frontal gyrus also demonstrated one of the biggest shifts between DBS-OFF and DBS-ON condition measuring fMRI (Saenger et al., 2017). Interestingly, the monophasic stimulus applied on STN provoked a slight decrease of activity in the supplementary motor area in our results. This finding is in line with experimental results showing that DBS weakens excessive phase-locking interactions in the motor areas of PD patients (de Hemptinne et al., 2015). Supplementary motor areas, which are located at the transition between primary motor areas and prefrontal cortex, are involved in intentional movement initiation (Goldberg, 1985) and their impaired function is supposed to contribute to PD symptoms (Jacobs et al., 2009). Direct stimulation of supplementary motor areas with transcranial magnetic stimulation leads to improved freezing of gait symptoms in PD (Kim et al., 2018; Shirota et al., 2013), while dopaminergic medication can be related to improved supplementary motor area activation and improved motoric functions (Jenkins et al., 1992; Rascol et al., 1994). STN DBS in PD has been shown in fMRI (Stefurak et al., 2003) and positron emission tomography (Ceballos-Baumann et al., 1999) studies to activate motor as well as premotor areas, concordant with the simulated patterns in this work.

Our spiking network relies on a high level of biological realism with regards to spatio-temporal dynamics. Space refers to the fact that the spiking network receives input from different brain regions of TVB, which is closer to the reality regarding the multitude of different white matter connections between the cortex and the BG (Lenglet et al., 2012). More realistic time modeling implies the specific mean-field model dynamics that are chosen, as opposed to other studies, in which spiking networks are driven by Poisson spike trains, white noise or harmonic oscillations (Humphries et al., 2006; Park et al., 2011; Terman et al., 2002). The former approach of driving these spiking networks with noise seems to be an abstract view of the biologically underlying phenomena (Kerr et al., 2013).

There are still several open challenges in the field of DBS research that a virtual testing environment could potentially address. First, the exact placement of the electrodes seems crucial for the clinical outcome for patients. For PD and OCD, recent studies have shown that the connectivity profile of the brain area encompassing the inserted electrode predicts clinical outcome measures for patients (Baldermann et al., 2019; Horn et al., 2017, 2019; Joutsa et al., 2018). This phenomenon was validated for dystonia (Corp et al., 2019; Okromelidze et al., 2020), essential tremor (Al-Fatly et al., 2019) and epilepsy (Middlebrooks et al., 2018). Testing the effects of different placement strategies before surgery could provide simulation-based advice for neurosurgeons. In this first co-simulation approach for DBS, we modeled stimuli targeting the GPi or STN area directly and completely. The clinical reality looks more complex (Krauss et al., 2021). The different sub-areas within the STN, for example, are involved in different pathways, i.e., the sensorimotor, associative and limbic loop. As most DBS systems provide several lead contacts to choose, the precise stimulus location is a common problem in clinical fine-tuning of DBS. With the upcoming of more detailed brain atlases, one could easily extend our used parcellation towards a finer grid inside the BG and model these subparts separately. Here, we presented the scaffold model that can be fine-tuned towards a more realistic model in a straight-forward manner. Second, so far, little individual information is considered for each patient and often the electrodes are placed based on normative data (Horn & Fox, 2020). Fitting an individual TVB model for patients provides a more personalized approach based on individual structural and functional imaging or electrophysiological data. TVB has previously been applied to help with predictions of clinical features for individual patients. Using individual positron emission tomography images, EEG slowing in patients with AD could be inferred from *Abeta* accumulation with the help of TVB (Stefanovski et al., 2019). Recently, a study has shown that the TVB feature of simulated mean local field potential frequency per brain region significantly improves the classification of individuals as AD patients, mild cognitive impairment patients or healthy controls using machine learning (Triebkorn et al., 2021). For epilepsy patients, TVB has successfully been applied to optimize the determination of the resection and epileptic zone per individual before surgery (An et al., 2019).

A personalized virtual brain including structural data and dynamics based on MRI data is flexible in exploring other neuromodulation techniques with little extra effort. The hypothesis is that neuromodulation techniques can move the brain network dynamics between the diseased and healthy state (Figure 10). With the current study, we have made a first attempt to “control” brain network dynamics by modeling stimulation in the brain of a PD patient. There is evidence that PD patients could also benefit from other neuromodulation techniques (Brittain & Cagnan, 2018). For example, a first study found that transcranial magnetic stimulation (TMS) of the supplementary motor area helps to improve the motoric symptoms of PD patients (Shirota et al., 2013). With our co-simulation framework, we can potentially analyze the impact of such a stimulation originating on the surface and follow the complete loop of cortico-basal-ganglia-thalamic-cortex connectivity. The flexibility of the presented virtual model could help with finding the best therapy for each individual patient.

**Figure 10:**
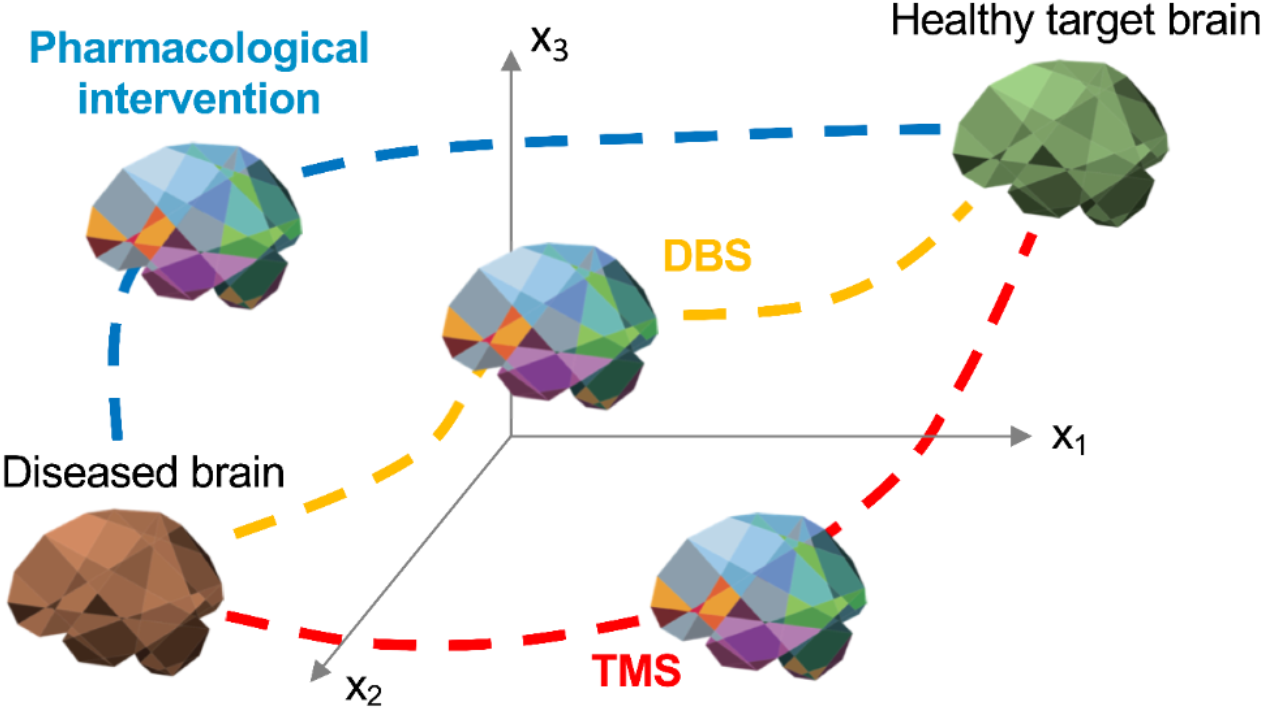
Schematic overview of “controlling” brain dynamics from one state to another using The Virtual Brain. Different interventions, e.g., deep brain stimulation (DBS), transcranial magnetic stimulation (TMS) or pharmacological interventions, can shift the brain dynamics from one state to another. These neuromodulation techniques hold the potential to alter the brain dynamics from a diseased brain towards a healthy target brain. Using The Virtual Brain, we aim to explore the different pathways leading to healthy functioning in a virtual environment for individual patients.

The applied data-driven model from Maith et al. (2020) does not make use of any prior assumptions regarding the pathological PD activity within the BG network, which stands in contrast to many previous models (Leblois et al., 2006; Lindahl & Hellgren Kotaleski, 2016). Fitting the outcomes of a model with empirical data from patients and controls offers an alternative approach to determining BG and whole-brain model dynamics. With this primarily data-driven approach, Maith et al. (2020) found many similarities of the obtained personal models of individual PD patients with physiological findings of PD, such as lower firing rates in the thalamus.

Our study inherits some limitations. In this proof-of-concept study, we modeled the TVB input that drives the spiking BG network with the reduced Wong-Wang mean-field model (Deco et al., 2013). In an improved version of this model, we could adjust the TVB mean-field dynamics to qualitatively correspond better with the original spiking network of Izhikevich neurons (Maith et al., 2020) by taking advantage of existing mean-field approximations of such networks (Nicola & Campbell, 2013; Visser & Van Gils, 2014). Such a choice would allow a more accurate analytical and computational determination of the large-scale brain dynamics (e.g., involved bifurcations) and inform the interface modeling between the two scales accordingly (e.g., in terms of scaling or more complex transformations). Further, alternatives to correlated spike trains’ generators for converting the TVB mean-field nodes’ rates into spike trains of TVB “proxy” nodes could be more effective in mimicking the Izhikevich spiking cortex node dynamics. All of the above options can be better explored by an upcoming computationally optimized version of the TVB-multiscale toolbox, implementing parallel co-simulation, allowing for a systematic exploration of the parameter space of the multiscale model to better fit individual neuroimaging data. So far, we fitted the virtual co-simulation brains to two individuals, which can be easily extended to larger cohorts with the only necessary data being DTI and either fMRI or electrophysiological data to fit the model dynamics accurately. The well-known characteristic of PD patients to demonstrate hyper-synchronization in the beta band (8-35*Hz*) in the sensorimotor network and the STN (Cruz et al., 2011; Whitmer et al., 2012) is reversed by DBS (Kühn et al., 2008; Wingeier et al., 2006). Our approach did not yet incorporate modeling the electrophysiological signatures of virtual DBS. However, TVB has often been used to monitor EEG-like activity from simulated time series and TVB-multiscale will soon also be equipped for this monitoring. Short-term plasticity probably plays an essential role in DBS effects (Milosevic et al., 2018), which has not yet been implemented in our model. Similarly, long-term plasticity effects due to DBS probably exist in structural and functional networks (van Hartevelt et al., 2014). With the spiking model allowing for an implementation of plasticity rules, we could explore its effects on the whole-brain dynamics with our model in future work. Moreover, our BG network misses the substantia nigra region as a crucial factor influencing PD dynamics and so far, we limited our analyses to a single (left) hemisphere.

## Conclusions

In this study, we presented a co-simulation model for the BG as a spiking network together with TVB mean-field simulations for the whole brain. Our results show biologically plausible effects of virtual DBS performed in this multiscale modeling framework, bringing the patient’s network dynamics of the BG closer to the healthy regime. The presented model offers a bridge between the different scales affected by DBS in the brain. It has the potential to be used as a ‘sandbox’ model for individual patients suffering from different neurological disorders prior to surgical interventions. Different strategies for DBS lead placements and configurations can be tested and evaluated. Future work needs to validate this model in larger patient cohorts and establish its link with clinical post-surgery improvement.

## Acknowledgments

PR acknowledges support by EU H2020 Virtual Brain Cloud 826421, Human Brain Project SGA2 785907; Human Brain Project SGA3 945539, ERC Consolidator 683049; German Research Foundation SFB 1436 (project ID 425899996); SFB 1315 (project ID 327654276); SFB 936 (project ID 178316478; SFB-TRR 295 (project ID 424778381); SPP Computational Connectomics RI 2073/6-1, RI 2073/10-2, RI 2073/9-1; Berlin Institute of Health & Foundation Charité, Johanna Quandt Excellence Initiative. In addition, we acknowledge support from the German Research Foundation (DFG) HA2630/11-1 and HA2630/11-2 part of “Computational Connectomics” (SPP 2041) and the Open Access Publication Fund of Charité-Universitätsmedizin Berlin. We thank Ningfei Li, Simón Oxenford and Andreas Horn for helping us prepare the connectome data for TVB and being involved in initial discussions. We are grateful to Julien Vitay for providing us with guidance concerning the usage of ANNarchy and being involved in initial discussions. We thank Arjan Hillebrand for sharing his code to visualize the template brains with us.

## References

Ackermans, L., Duits, A., van der Linden, C., Tijssen, M., Schruers, K., Temel, Y., Kleijer, M., Nederveen, P., Bruggeman, R., Tromp, S., van Kranen-Mastenbroek, V., Kingma, H., Cath, D., & Visser-Vandewalle, V. (2011). Double-blind clinical trial of thalamic stimulation in patients with Tourette syndrome. Brain: A Journal of Neurology, 134(Pt 3), 832–844. https://doi.org/10.1093/brain/awq380

Aerts, H., Schirner, M., Jeurissen, B., Van Roost, D., Achten, E., Ritter, P., & Marinazzo, D. (2018). Modeling Brain Dynamics in Brain Tumor Patients Using the Virtual Brain. eNeuro, 5(3). https://doi.org/10.1523/ENEURO.0083-18.2018

Albin, R. L., Young, A. B., & Penney, J. B. (1989). The functional anatomy of basal ganglia disorders. Trends in Neurosciences, 12(10), 366–375. https://doi.org/10.1016/0166-2236(89)90074-x

Al-Fatly, B., Ewert, S., Kübler, D., Kroneberg, D., Horn, A., & Kühn, A. A. (2019). Connectivity profile of thalamic deep brain stimulation to effectively treat essential tremor. Brain: A Journal of Neurology, 142(10), 3086–3098. https://doi.org/10.1093/brain/awz236

Anderson, D., & Ahmed, A. (2003). Treatment of patients with intractable obsessive-compulsive disorder with anterior capsular stimulation. Case report. Journal of Neurosurgery, 98(5), 1104–1108. https://doi.org/10.3171/jns.2003.98.5.1104

An, S., Bartolomei, F., Guye, M., & Jirsa, V. (2019). Optimization of surgical intervention outside the epileptogenic zone in the Virtual Epileptic Patient (VEP). PLoS Computational Biology, 15(6), e1007051. https://doi.org/10.1371/journal.pcbi.1007051

Baladron, J., & Hamker, F. H. (2020). Habit learning in hierarchical cortex-basal ganglia loops. The European Journal of Neuroscience, 52(12), 4613–4638. https://doi.org/10.1111/ejn.14730

Baladron, J., Nambu, A., & Hamker, F. H. (2019). The subthalamic nucleus-external globus pallidus loop biases exploratory decisions towards known alternatives: a neuro-computational study. The European Journal of Neuroscience, 49(6), 754–767. https://doi.org/10.1111/ejn.13666

Baldermann, J. C., Melzer, C., Zapf, A., Kohl, S., Timmermann, L., Tittgemeyer, M., Huys, D., Visser-Vandewalle, V., Kühn, A. A., Horn, A., & Kuhn, J. (2019). Connectivity Profile Predictive of Effective Deep Brain Stimulation in Obsessive-Compulsive Disorder. Biological Psychiatry, 85(9), 735–743. https://doi.org/10.1016/j.biopsych.2018.12.019

Bergelt, J., & Hamker, F. H. (2019). Spatial updating of attention across eye movements: A neuro-computational approach. Journal of Vision, 19(7), 10. https://doi.org/10.1167/19.7.10

Brittain, J.-S., & Cagnan, H. (2018). Recent Trends in the Use of Electrical Neuromodulation in Parkinson’s Disease. Current Behavioral Neuroscience Reports, 5(2), 170–178. https://doi.org/10.1007/s40473-018-0154-9

Ceballos-Baumann, A. O., Boecker, H., Bartenstein, P., von Falkenhayn, I., Riescher, H., Conrad, B., Moringlane, J. R., & Alesch, F. (1999). A positron emission tomographic study of subthalamic nucleus stimulation in Parkinson disease: enhanced movement-related activity of motor-association cortex and decreased motor cortex resting activity. Archives of Neurology, 56(8), 997–1003. https://doi.org/10.1001/archneur.56.8.997

Chiken, S., & Nambu, A. (2016). Mechanism of Deep Brain Stimulation: Inhibition, Excitation, or Disruption? The Neuroscientist: A Review Journal Bringing Neurobiology, Neurology and Psychiatry, 22(3), 313–322. https://doi.org/10.1177/1073858415581986

Christopher, L., Koshimori, Y., Lang, A. E., Criaud, M., & Strafella, A. P. (2014). Uncovering the role of the insula in non-motor symptoms of Parkinson’s disease. In Brain (Vol. 137, Issue 8, pp. 2143–2154). https://doi.org/10.1093/brain/awu084

Corp, D. T., Joutsa, J., Darby, R. R., Delnooz, C. C. S., van de Warrenburg, B. P. C., Cooke, D., Prudente, C. N., Ren, J., Reich, M. M., Batla, A., Bhatia, K. P., Jinnah, H. A., Liu, H., & Fox, M. D. (2019). Network localization of cervical dystonia based on causal brain lesions. Brain: A Journal of Neurology, 142(6), 1660–1674. https://doi.org/10.1093/brain/awz112

Cruz, A. V., Mallet, N., Magill, P. J., Brown, P., & Averbeck, B. B. (2011). Effects of dopamine depletion on information flow between the subthalamic nucleus and external globus pallidus. Journal of Neurophysiology, 106(4), 2012–2023. https://doi.org/10.1152/jn.00094.2011

Damier, P., Hirsch, E. C., Agid, Y., & Graybiel, A. M. (1999). The substantia nigra of the human brain. II. Patterns of loss of dopamine-containing neurons in Parkinson’s disease. Brain: A Journal of Neurology, 122 (Pt 8), 1437–1448. https://doi.org/10.1093/brain/122.8.1437

Deco, G., Ponce-Alvarez, A., Mantini, D., Romani, G. L., Hagmann, P., & Corbetta, M. (2013). Resting-State Functional Connectivity Emerges from Structurally and Dynamically Shaped Slow Linear Fluctuations. The Journal of Neuroscience: The Official Journal of the Society for Neuroscience, 33(27), 11239–11252. https://doi.org/10.1523/JNEUROSCI.1091-13.2013

de Hemptinne, C., Swann, N. C., Ostrem, J. L., Ryapolova-Webb, E. S., San Luciano, M., Galifianakis, N. B., & Starr, P. A. (2015). Therapeutic deep brain stimulation reduces cortical phase-amplitude coupling in Parkinson’s disease. Nature Neuroscience, 18(5), 779–786. https://doi.org/10.1038/nn.3997

DeLong, M. R. (1990). Primate models of movement disorders of basal ganglia origin. Trends in Neurosciences, 13(7), 281–285. https://doi.org/10.1016/0166-2236(90)90110-v

Deuschl, G., Schade-Brittinger, C., Krack, P., Volkmann, J., Schäfer, H., Bötzel, K., Daniels, C., Deutschländer, A., Dillmann, U., Eisner, W., Gruber, D., Hamel, W., Herzog, J., Hilker, R., Klebe, S., Kloß, M., Koy, J., Krause, M., Kupsch, A.,… Voges, J. (2006). A Randomized Trial of Deep-Brain Stimulation for Parkinson’s Disease. In New England Journal of Medicine (Vol. 355, Issue 9, pp. 896–908). https://doi.org/10.1056/nejmoa060281

Dostrovsky, J. O., Hutchison, W. D., & Lozano, A. M. (2002). The Globus Pallidus, Deep Brain Stimulation, and Parkinson’s Disease. In The Neuroscientist (Vol. 8, Issue 3, pp. 284–290). https://doi.org/10.1177/1073858402008003014

Eisinger, R. S., Cernera, S., Gittis, A., Gunduz, A., & Okun, M. S. (2019). A review of basal ganglia circuits and physiology: application to deep brain stimulation. Parkinsonism & Related Disorders, 59, 9. https://doi.org/10.1016/j.parkreldis.2019.01.009

Eppler, J. M., Helias, M., Muller, E., Diesmann, M., & Gewaltig, M.-O. (2008). PyNEST: A Convenient Interface to the NEST Simulator. Frontiers in Neuroinformatics, 2, 12. https://doi.org/10.3389/neuro.11.012.2008

Ewert, S., Plettig, P., Li, N., Chakravarty, M. M., Collins, D. L., Herrington, T. M., Kühn, A. A., & Horn, A. (2018). Toward defining deep brain stimulation targets in MNI space: A subcortical atlas based on multimodal MRI, histology and structural connectivity. NeuroImage, 170, 271–282. https://doi.org/10.1016/j.neuroimage.2017.05.015

Fearnley, J. M., & Lees, A. J. (1991). Ageing and Parkinson’s disease: substantia nigra regional selectivity. Brain: A Journal of Neurology, 114 (Pt 5), 2283–2301. https://doi.org/10.1093/brain/114.5.2283

Franzini, A., Messina, G., Gambini, O., Muffatti, R., Scarone, S., Cordella, R., & Broggi, G. (2010). Deep-brain stimulation of the nucleus accumbens in obsessive compulsive disorder: clinical, surgical and electrophysiological considerations in two consecutive patients. Neurological Sciences: Official Journal of the Italian Neurological Society and of the Italian Society of Clinical Neurophysiology, 31(3), 353–359. https://doi.org/10.1007/s10072-009-0214-8

Goldberg, G. (1985). Supplementary motor area structure and function: Review and hypotheses. In Behavioral and Brain Sciences (Vol. 8, Issue 4, pp. 567–588). https://doi.org/10.1017/s0140525x00045167

Gönner, L., Maith, O., Koulouri, I., Baladron, J., & Hamker, F. H. (2020). A spiking model of basal ganglia dynamics in stopping behavior supported by arkypallidal neurons. The European Journal of Neuroscience. https://doi.org/10.1111/ejn.15082

Gönner, L., Vitay, J., & Hamker, F. H. (2017). Predictive Place-Cell Sequences for Goal-Finding Emerge from Goal Memory and the Cognitive Map: A Computational Model. Frontiers in Computational Neuroscience, 11, 84. https://doi.org/10.3389/fncom.2017.00084

Gruber, D., Kuhn, A. A., Schoenecker, T., Kopp, U. A., Kivi, A., Huebl, J., Lobsien, E., Mueller, B., Schneider, G.-H., & Kupsch, A. (2014). Quadruple deep brain stimulation in Huntington’s disease, targeting pallidum and subthalamic nucleus: case report and review of the literature. Journal of Neural Transmission, 121(10), 1303–1312. https://doi.org/10.1007/s00702-014-1201-7

Guo, Y., Rubin, J. E., McIntyre, C. C., Vitek, J. L., & Terman, D. (2008). Thalamocortical Relay Fidelity Varies Across Subthalamic Nucleus Deep Brain Stimulation Protocols in a Data-Driven Computational Model. In Journal of Neurophysiology (Vol. 99, Issue 3, pp. 1477–1492). https://doi.org/10.1152/jn.01080.2007

Hahn, P. J., & McIntyre, C. C. (2010). Modeling shifts in the rate and pattern of subthalamopallidal network activity during deep brain stimulation. In Journal of Computational Neuroscience (Vol. 28, Issue 3, pp. 425–441). https://doi.org/10.1007/s10827-010-0225-8

Hahn, P. J., Russo, G. S., Hashimoto, T., Miocinovic, S., Xu, W., McIntyre, C. C., & Vitek, J. L. (2008). Pallidal burst activity during therapeutic deep brain stimulation. Experimental Neurology, 211(1), 243–251. https://doi.org/10.1016/j.expneurol.2008.01.032

Hashimoto, T., Elder, C. M., Okun, M. S., Patrick, S. K., & Vitek, J. L. (2003). Stimulation of the subthalamic nucleus changes the firing pattern of pallidal neurons. The Journal of Neuroscience: The Official Journal of the Society for Neuroscience, 23(5), 1916–1923. https://www.ncbi.nlm.nih.gov/pubmed/12629196

Heimer, L. (1983). Basal Ganglia. In The Human Brain and Spinal Cord (pp. 199–209). Springer, New York, NY. https://doi.org/10.1007/978-1-4684-0150-9_15

Horn, A. (2019). The impact of modern-day neuroimaging on the field of deep brain stimulation. Current Opinion in Neurology, 32(4), 511–520. https://doi.org/10.1097/WCO.0000000000000679

Horn, A., & Fox, M. D. (2020). Opportunities of connectomic neuromodulation. NeuroImage, 221, 117180. https://doi.org/10.1016/j.neuroimage.2020.117180

Horn, A., Reich, M., Vorwerk, J., Li, N., Wenzel, G., Fang, Q., Schmitz-Hübsch, T., Nickl, R., Kupsch, A., Volkmann, J., Kühn, A. A., & Fox, M. D. (2017). Connectivity Predicts deep brain stimulation outcome in Parkinson disease. In Annals of Neurology (Vol. 82, Issue 1, pp. 67–78). https://doi.org/10.1002/ana.24974

Horn, A., Wenzel, G., Irmen, F., Huebl, J., Li, N., Neumann, W.-J., Krause, P., Bohner, G., Scheel, M., & Kühn, A. A. (2019). Deep brain stimulation induced normalization of the human functional connectome in Parkinson’s disease. Brain: A Journal of Neurology, 142(10), 3129–3143. https://doi.org/10.1093/brain/awz239

Humphries, M. D., & Gurney, K. (2012). Network effects of subthalamic deep brain stimulation drive a unique mixture of responses in basal ganglia output. In European Journal of Neuroscience (Vol. 36, Issue 2, pp. 2240–2251). https://doi.org/10.1111/j.1460-9568.2012.08085.x

Humphries, M. D., Obeso, J. A., & Dreyer, J. K. (2018). Insights into Parkinson’s disease from computational models of the basal ganglia. Journal of Neurology, Neurosurgery, and Psychiatry, 89(11), 1181–1188. https://doi.org/10.1136/jnnp-2017-315922

Humphries, M. D., Stewart, R. D., & Gurney, K. N. (2006). A physiologically plausible model of action selection and oscillatory activity in the basal ganglia. The Journal of Neuroscience: The Official Journal of the Society for Neuroscience, 26(50), 12921– 12942. https://doi.org/10.1523/JNEUROSCI.3486-06.2006

Irmen, F., Horn, A., Meder, D., Neumann, W.-J., Plettig, P., Schneider, G.-H., Siebner, H. R., & Kühn, A. A. (2019). Sensorimotor subthalamic stimulation restores risk-reward trade-off in Parkinson’s disease. Movement Disorders: Official Journal of the Movement Disorder Society, 34(3), 366–376. https://doi.org/10.1002/mds.27576

Izhikevich, E. M. (2004). Which model to use for cortical spiking neurons? IEEE Transactions on Neural Networks / a Publication of the IEEE Neural Networks Council, 15(5), 1063–1070. https://doi.org/10.1109/TNN.2004.832719

Jacobs, J. V., Lou, J. S., Kraakevik, J. A., & Horak, F. B. (2009). The supplementary motor area contributes to the timing of the anticipatory postural adjustment during step initiation in participants with and without Parkinson’s disease. Neuroscience, 164(2), 877–885. https://doi.org/10.1016/j.neuroscience.2009.08.002

Jahanshahi, M., Obeso, I., Baunez, C., Alegre, M., & Krack, P. (2015). Parkinson’s disease, the subthalamic nucleus, inhibition, and impulsivity. Movement Disorders: Official Journal of the Movement Disorder Society, 30(2), 128–140. https://doi.org/10.1002/mds.26049

Jamalian, A., Bergelt, J.,Dinkelbach, H.Ü., & Hamker, F. H. (2017). Spatial Attention Improves Object Localization: A Biologically Plausible Neuro-Computational Model for Use in Virtual Reality. 2017 IEEE International Conference on Computer Vision Workshops (ICCVW), 2724–2729. https://doi.org/10.1109/ICCVW.2017.320

Jenkins, I. H., Fernandez, W., Playford, E. D., Lees, A. J., Frackowiak, R. S., Passingham, R. E., & Brooks, D. J. (1992). Impaired activation of the supplementary motor area in Parkinson’s disease is reversed when akinesia is treated with apomorphine. Annals of Neurology, 32(6), 749–757. https://doi.org/10.1002/ana.410320608

Jones, D. K., Knösche, T. R., & Turner, R. (2013). White matter integrity, fiber count, and other fallacies: the do’s and don’ts of diffusion MRI. NeuroImage, 73, 239–254. https://doi.org/10.1016/j.neuroimage.2012.06.081

Joutsa, J., Horn, A., Hsu, J., & Fox, M. D. (2018). Localizing parkinsonism based on focal brain lesions. In Brain (Vol. 141, Issue 8, pp. 2445–2456). https://doi.org/10.1093/brain/awy161

Kahan, J., Mancini, L., Urner, M., Friston, K., Hariz, M., Holl, E., White, M., Ruge, D., Jahanshahi, M., Boertien, T., Yousry, T., Thornton, J. S., Limousin, P., Zrinzo, L., & Foltynie, T. (2012). Therapeutic subthalamic nucleus deep brain stimulation reverses cortico-thalamic coupling during voluntary movements in Parkinson’s disease. PloS One, 7(12), e50270. https://doi.org/10.1371/journal.pone.0050270

Kerr, C. C., Van Albada, S. J., Neymotin, S. A., Chadderdon, G. L., Robinson, P. A., & Lytton, W. W. (2013). Cortical information flow in Parkinson’s disease: a composite network/field model. Frontiers in Computational Neuroscience, 7, 39. https://doi.org/10.3389/fncom.2013.00039

Kim, S. J., Paeng, S. H., & Kang, S. Y. (2018). Stimulation in Supplementary Motor Area Versus Motor Cortex for Freezing of Gait in Parkinson’s Disease. Journal of Clinical Neurology, 14(3), 320–326. https://doi.org/10.3988/jcn.2018.14.3.320

Klein, P. C., Ettinger, U., Schirner, M., Ritter, P., Rujescu, D., Falkai, P., Koutsouleris, N., Kambeitz-Ilankovic, L., & Kambeitz, J. (2021). Brain Network Simulations Indicate Effects of Neuregulin-1 Genotype on Excitation-Inhibition Balance in Cortical Dynamics. Cerebral Cortex, 31(4), 2013–2025. https://doi.org/10.1093/cercor/bhaa339

Koller, W., Pahwa, R., Busenbark, K., Hubble, J., Wilkinson, S., Lang, A., Tuite, P., Sime, E., Lazano, A., Hauser, R., Malapira, T., Smith, D., Tarsy, D., Miyawaki, E., Norregaard, T., Kormos, T., & Olanow, C. W. (1997). High-frequency unilateral thalamic stimulation in the treatment of essential and parkinsonian tremor. Annals of Neurology, 42(3), 292– 299. https://doi.org/10.1002/ana.410420304

Krauss, J. K., Lipsman, N., Aziz, T., Boutet, A., Brown, P., Chang, J. W., Davidson, B., Grill, W. M., Hariz, M. I., Horn, A., Schulder, M., Mammis, A., Tass, P. A., Volkmann, J., & Lozano, A. M. (2021). Technology of deep brain stimulation: current status and future directions. Nature Reviews. Neurology, 17(2), 75–87. https://doi.org/10.1038/s41582-020-00426-z

Kühn, A. A., Kempf, F., Brücke, C., Gaynor Doyle, L., Martinez-Torres, I., Pogosyan, A., Trottenberg, T., Kupsch, A., Schneider, G.-H., Hariz, M. I., Vandenberghe, W., Nuttin, B., & Brown, P. (2008). High-frequency stimulation of the subthalamic nucleus suppresses oscillatory beta activity in patients with Parkinson’s disease in parallel with improvement in motor performance. The Journal of Neuroscience: The Official Journal of the Society for Neuroscience, 28(24), 6165–6173. https://doi.org/10.1523/JNEUROSCI.0282-08.2008

Kunze, T., Hunold, A., Haueisen, J., Jirsa, V., & Spiegler, A. (2016). Transcranial direct current stimulation changes resting state functional connectivity: A large-scale brain network modeling study. NeuroImage, 140, 174–187. https://doi.org/10.1016/j.neuroimage.2016.02.015

Kupsch, A., Benecke, R., Müller, J., Trottenberg, T., Schneider, G.-H., Poewe, W., Eisner, W., Wolters, A., Müller, J.-U., Deuschl, G., Pinsker, M. O., Skogseid, I. M., Roeste, G. K., Vollmer-Haase, J., Brentrup, A., Krause, M., Tronnier, V., Schnitzler, A., Voges, J.,… Volkmann, J. (2006). Pallidal Deep-Brain Stimulation in Primary Generalized or Segmental Dystonia. In New England Journal of Medicine (Vol. 355, Issue 19, pp. 1978–1990). https://doi.org/10.1056/nejmoa063618

Lanciego, J. L., Luquin, N., & Obeso, J. A. (2012). Functional Neuroanatomy of the Basal Ganglia. Cold Spring Harbor Perspectives in Medicine, 2(12). https://doi.org/10.1101/cshperspect.a009621

Larisch, R., Gönner, L., Teichmann, M., & Hamker, F. H. (2020). Sensory coding and contrast invariance emerge from the control of plastic inhibition over excitatory connectivity. In Cold Spring Harbor Laboratory (p. 2020.04.07.029157). https://doi.org/10.1101/2020.04.07.029157

Leblois, A., Boraud, T., Meissner, W., Bergman, H., & Hansel, D. (2006). Competition between feedback loops underlies normal and pathological dynamics in the basal ganglia. The Journal of Neuroscience: The Official Journal of the Society for Neuroscience, 26(13), 3567–3583. https://doi.org/10.1523/JNEUROSCI.5050-05.2006

Lenglet, C., Abosch, A., Yacoub, E., De Martino, F., Sapiro, G., & Harel, N. (2012). Comprehensive in vivo mapping of the human basal ganglia and thalamic connectome in individuals using 7T MRI. PloS One, 7(1), e29153. https://doi.org/10.1371/journal.pone.0029153

Lindahl, M., &Hellgren Kotaleski, J. (2016). Untangling Basal Ganglia Network Dynamics and Function: Role of Dopamine Depletion and Inhibition Investigated in a Spiking Network Model. eNeuro, 3(6). https://doi.org/10.1523/ENEURO.0156-16.2016

Liu, C., Zhao, G., Wang, J., Wu, H., Li, H., Fietkiewicz, C., & Loparo, K. A. (2020). Neural network-based closed-loop deep brain stimulation for modulation of pathological oscillation in Parkinson’s disease. IEEE Access, 8, 161067–161079. https://ieeexplore.ieee.org/abstract/document/9180308/

Maith, O., Villagrasa Escudero, F., Dinkelbach, H.Ü., Baladron, J., Horn, A., Irmen, F., Kühn, A. A., & Hamker, F. H. (2020). A computational model-based analysis of basal ganglia pathway changes in Parkinson’s disease inferred from resting-state fMRI. The European Journal of Neuroscience. https://doi.org/10.1111/ejn.14868

Marsden, C. D., & Obeso, J. A. (1994). The functions of the basal ganglia and the paradox of stereotaxic surgery in Parkinson’s disease. Brain: A Journal of Neurology, 117 (Pt 4), 877–897. https://doi.org/10.1093/brain/117.4.877

Mayberg, H. S., Lozano, A. M., Voon, V., McNeely, H. E., Seminowicz, D., Hamani, C., Schwalb, J. M., & Kennedy, S. H. (2005). Deep Brain Stimulation for Treatment-Resistant Depression. In Neuron (Vol. 45, Issue 5, pp. 651–660). https://doi.org/10.1016/j.neuron.2005.02.014

McIntyre, C. C., & Hahn, P. J. (2010). Network perspectives on the mechanisms of deep brain stimulation. In Neurobiology of Disease (Vol. 38, Issue 3, pp. 329–337). https://doi.org/10.1016/j.nbd.2009.09.022

Michmizos, K. P., & Nikita, K. S. (2011). Addition of deep brain stimulation signal to a local field potential driven Izhikevich model masks the pathological firing pattern of an STN neuron. Conference Proceedings: … Annual International Conference of the IEEE Engineering in Medicine and Biology Society. IEEE Engineering in Medicine and Biology Society. Conference, 2011, 7290–7293. https://doi.org/10.1109/IEMBS.2011.6091700

Middlebrooks, E. H., Grewal, S. S., Stead, M., Lundstrom, B. N., Worrell, G. A., & Van Gompel, J. J. (2018). Differences in functional connectivity profiles as a predictor of response to anterior thalamic nucleus deep brain stimulation for epilepsy: a hypothesis for the mechanism of action and a potential biomarker for outcomes. Neurosurgical Focus, 45(2), E7. https://doi.org/10.3171/2018.5.FOCUS18151

Milosevic, L., Kalia, S. K., Hodaie, M., Lozano, A. M., Fasano, A., Popovic, M. R., & Hutchison, W. D. (2018). Neuronal inhibition and synaptic plasticity of basal ganglia neurons in Parkinson’s disease. Brain: A Journal of Neurology, 141(1), 177–190. https://doi.org/10.1093/brain/awx296

Moran, A., Stein, E., Tischler, H., & Bar-Gad, I. (2012). Decoupling neuronal oscillations during subthalamic nucleus stimulation in the parkinsonian primate. In Neurobiology of Disease (Vol. 45, Issue 1, pp. 583–590). https://doi.org/10.1016/j.nbd.2011.09.016

Müller, J., Nawrot, M., Menzel, R., & Landgraf, T. (2018). A neural network model for familiarity and context learning during honeybee foraging flights. Biological Cybernetics, 112(1-2), 113–126. https://doi.org/10.1007/s00422-017-0732-z

Müller, U. J., Sturm, V., Voges, J., Heinze, H.-J., Galazky, I., Heldmann, M., Scheich, H., & Bogerts, B. (2009). Successful treatment of chronic resistant alcoholism by deep brain stimulation of nucleus accumbens: first experience with three cases. Pharmacopsychiatry, 42(6), 288–291. https://doi.org/10.1055/s-0029-1233489

Nambu, A., Tokuno, H., & Takada, M. (2002). Functional significance of the cortico– subthalamo–pallidal “hyperdirect” pathway. In Neuroscience Research (Vol. 43, Issue 2, pp. 111–117). https://doi.org/10.1016/s0168-0102(02)00027-5

Nicola, W., & Campbell, S. A. (2013). Mean-field models for heterogeneous networks of two-dimensional integrate and fire neurons. Frontiers in Computational Neuroscience, 7. https://doi.org/10.3389/fncom.2013.00184

Noecker, A. M., Frankemolle-Gilbert, A. M., Howell, B., Petersen, M. V., Beylergil, S. B., Shaikh, A. G., & McIntyre, C. C. (2021). StimVision v2: Examples and Applications in Subthalamic Deep Brain Stimulation for Parkinson’s Disease. Neuromodulation: Journal of the International Neuromodulation Society, 24(2), 248–258. https://doi.org/10.1111/ner.13350

Nuttin, B. J., Gabriëls, L. A., Cosyns, P. R., Meyerson, B. A., Andréewitch, S., Sunaert, S. G., Maes, A. F., Dupont, P. J., Gybels, J. M., Gielen, F., & Demeulemeester, H. G. (2008). Long-term electrical capsular stimulation in patients with obsessive-compulsive disorder. Neurosurgery, 62(6 Suppl 3), 966–977. https://doi.org/10.1227/01.neu.0000333764.20575.d6

Okromelidze, L., Tsuboi, T., Eisinger, R. S., Burns, M. R., Charbel, M., Rana, M., Grewal, S. S., Lu, C.-Q., Almeida, L., Foote, K. D., Okun, M. S., & Middlebrooks, E. H. (2020). Functional and Structural Connectivity Patterns Associated with Clinical Outcomes in Deep Brain Stimulation of the Globus Pallidus Internus for Generalized Dystonia. AJNR. American Journal of Neuroradiology, 41(3), 508–514. https://doi.org/10.3174/ajnr.A6429

Park, C., Worth, R. M., & Rubchinsky, L. L. (2011). Neural dynamics in parkinsonian brain: the boundary between synchronized and nonsynchronized dynamics. Physical Review. E, Statistical, Nonlinear, and Soft Matter Physics, 83(4 Pt 1), 042901. https://doi.org/10.1103/PhysRevE.83.042901

Pauli, W. M., Nili, A. N., & Tyszka, J. M. (2018). A high-resolution probabilistic in vivo atlas of human subcortical brain nuclei. Scientific Data, 5, 180063. https://doi.org/10.1038/sdata.2018.63

Petersen, M. V., Mlakar, J., Haber, S. N., Parent, M., Smith, Y., Strick, P. L., Griswold, M. A., & McIntyre, C. C. (2019). Holographic Reconstruction of Axonal Pathways in the Human Brain. Neuron, 104(6), 1056–1064.e3. https://doi.org/10.1016/j.neuron.2019.09.030

Plotkin, J. L., & Goldberg, J. A. (2019). Thinking Outside the Box (and Arrow): Current Themes in Striatal Dysfunction in Movement Disorders. The Neuroscientist: A Review Journal Bringing Neurobiology, Neurology and Psychiatry, 25(4), 359–379. https://doi.org/10.1177/1073858418807887

Rascol, O., Sabatini, U., Chollet, F., Fabre, N., Senard, J. M., Montastruc, J. L., Celsis, P., Marc-Vergnes, J. P., & Rascol, A. (1994). Normal activation of the supplementary motor area in patients with Parkinson’s disease undergoing long-term treatment with levodopa. Journal of Neurology, Neurosurgery, and Psychiatry, 57(5), 567–571. https://doi.org/10.1136/jnnp.57.5.567

Reese, R., Leblois, A., Steigerwald, F., Pötter-Nerger, M., Herzog, J., Maximilian Mehdorn, H., Deuschl, G., Meissner, W. G., & Volkmann, J. (2011). Subthalamic deep brain stimulation increases pallidal firing rate and regularity. In Experimental Neurology (Vol. 229, Issue 2, pp. 517–521). https://doi.org/10.1016/j.expneurol.2011.01.020

Ritter, P., Schirner, M., McIntosh, A. R., & Jirsa, V. K. (2013). The virtual brain integrates computational modeling and multimodal neuroimaging. Brain Connectivity, 3(2), 121– 145. https://doi.org/10.1089/brain.2012.0120

Rodriguez-Oroz, M. C., Jahanshahi, M., Krack, P., Litvan, I., Macias, R., Bezard, E., & Obeso, J. A. (2009). Initial clinical manifestations of Parkinson’s disease: features and pathophysiological mechanisms. Lancet Neurology, 8(12), 1128–1139. https://doi.org/10.1016/S1474-4422(09)70293-5

Rolls, E., Joliot, M., & Tzourio-Mazoyer, N. (2015). Implementation of a new parcellation of the orbitofrontal cortex in the automated anatomical labeling atlas. NeuroImage, 122, 1– 5. https://doi.org/10.1016/j.neuroimage.2015.07.075

Rosenbaum, R., Zimnik, A., Zheng, F., Turner, R. S., Alzheimer, C., Doiron, B., & Rubin, J. E. (2014). Axonal and synaptic failure suppress the transfer of firing rate oscillations, synchrony and information during high frequency deep brain stimulation. In Neurobiology of Disease (Vol. 62, pp. 86–99). https://doi.org/10.1016/j.nbd.2013.09.006

Rubin, J. E., McIntyre, C. C., Turner, R. S., & Wichmann, T. (2012). Basal ganglia activity patterns in parkinsonism and computational modeling of their downstream effects. In European Journal of Neuroscience (Vol. 36, Issue 2, pp. 2213–2228). https://doi.org/10.1111/j.1460-9568.2012.08108.x

Rubin, J. E., & Terman, D. (2004). High Frequency Stimulation of the Subthalamic Nucleus Eliminates Pathological Thalamic Rhythmicity in a Computational Model. In Journal of Computational Neuroscience (Vol. 16, Issue 3, pp. 211–235). https://doi.org/10.1023/b:jcns.0000025686.47117.67

Saenger, V. M., Kahan, J., Foltynie, T., Friston, K., Aziz, T. Z., Green, A. L., van Hartevelt, T. J., Cabral, J., Stevner, A. B. A., Fernandes, H. M., Mancini, L., Thornton, J., Yousry, T., Limousin, P., Zrinzo, L., Hariz, M., Marques, P., Sousa, N., Kringelbach, M. L., & Deco, G. (2017). Uncovering the underlying mechanisms and whole-brain dynamics of deep brain stimulation for Parkinson’s disease. Scientific Reports, 7(1), 9882. https://doi.org/10.1038/s41598-017-10003-y

Salanova, V., Witt, T., Worth, R., Henry, T. R., Gross, R. E., Nazzaro, J. M., Labar, D., Sperling, M. R., Sharan, A., Sandok, E., Handforth, A., Stern, J. M., Chung, S., Henderson, J. M., French, J., Baltuch, G., Rosenfeld, W. E., Garcia, P., Barbaro, N. M.,… SANTE Study Group. (2015). Long-term efficacy and safety of thalamic stimulation for drug-resistant partial epilepsy. Neurology, 84(10), 1017–1025. https://doi.org/10.1212/WNL.0000000000001334

Sanz Leon, P., Knock, S. A., Woodman, M. M., Domide, L., Mersmann, J., McIntosh, A. R., & Jirsa, V. (2013). The Virtual Brain: a simulator of primate brain network dynamics. Frontiers in Neuroinformatics, 7, 10. https://doi.org/10.3389/fninf.2013.00010

Schirner, M., Domide, L., Perdikis, D., Triebkorn, P., Stefanovski, L., Pai, R., Popa, P., Valean, B., Palmer, J., Langford, C., Blickensdörfer, A., van der Vlag, M., Diaz-Pier, S., Peyser, A., Klijn, W., Pleiter, D., Nahm, A., Schmid, O., Woodman, M.,… Ritter, P. (2021). Brain Modelling as a Service: The Virtual Brain on EBRAINS. In arXiv [cs.CE]. arXiv. http://arxiv.org/abs/2102.05888

Schirner, M., McIntosh, A. R., Jirsa, V., Deco, G., & Ritter, P. (2018). Inferring multi-scale neural mechanisms with brain network modelling. eLife, 7. https://doi.org/10.7554/eLife.28927

Schmid, K., Vitay, J., & Hamker, F. H. (2019). Forward models in the cerebellum using reservoirs and perturbation learning. 2019 Conference on Cognitive Computational Neuroscience. 2019 Conference on Cognitive Computational Neuroscience, Berlin, Germany. https://doi.org/10.32470/ccn.2019.1139-0

Shirota, Y., Ohtsu, H., Hamada, M., Enomoto, H., Ugawa, Y., & For the Research Committee on rTMS Treatment of Parkinson’s Disease. (2013). Supplementary motor area stimulation for Parkinson disease: A randomized controlled study. In Neurology (Vol. 80, Issue 15, pp. 1400–1405). https://doi.org/10.1212/wnl.0b013e31828c2f66

Spiegler, A., Abadchi, J. K., Mohajerani, M., & Jirsa, V. K. (2020). In silico exploration of mouse brain dynamics by focal stimulation reflects the organization of functional networks and sensory processing. In Network Neuroscience (Vol. 4, Issue 3, pp. 807– 851). https://doi.org/10.1162/netn_a_00152

Spiegler, A., Hansen, E. C. A., Bernard, C., McIntosh, A. R., & Jirsa, V. K. (2016). Selective Activation of Resting-State Networks following Focal Stimulation in a Connectome-Based Network Model of the Human Brain. eNeuro, 3(5). https://doi.org/10.1523/ENEURO.0068-16.2016

Stefanovski, L., Triebkorn, P., Spiegler, A., Diaz-Cortes, M.-A., Solodkin, A., Jirsa, V., McIntosh, A. R., Ritter, P., & Alzheimer’s Disease Neuroimaging Initiative. (2019). Linking Molecular Pathways and Large-Scale Computational Modeling to Assess Candidate Disease Mechanisms and Pharmacodynamics in Alzheimer’s Disease. Frontiers in Computational Neuroscience, 13, 54. https://doi.org/10.3389/fncom.2019.00054

Stefurak, T., Mikulis, D., Mayberg, H., Lang, A. E., Hevenor, S., Pahapill, P., Saint-Cyr, J., & Lozano, A. (2003). Deep brain stimulation for Parkinson’s disease dissociates mood and motor circuits: a functional MRI case study. Movement Disorders: Official Journal of the Movement Disorder Society, 18(12), 1508–1516. https://doi.org/10.1002/mds.10593

Terman, D., Rubin, J. E., Yew, A. C., & Wilson, C. J. (2002). Activity Patterns in a Model for the Subthalamopallidal Network of the Basal Ganglia. In The Journal of Neuroscience (Vol. 22, Issue 7, pp. 2963–2976). https://doi.org/10.1523/jneurosci.22-07-02963.2002

Thomas, C., Ye, F. Q., Irfanoglu, M. O., Modi, P., Saleem, K. S., Leopold, D. A., & Pierpaoli, C. (2014). Anatomical accuracy of brain connections derived from diffusion MRI tractography is inherently limited. Proceedings of the National Academy of Sciences of the United States of America, 111(46), 16574–16579. https://doi.org/10.1073/pnas.1405672111

Triebkorn, P., Stefanovski, L., Dhindsa, K., Diaz-Cortes, M.-A., Bey, P., Bülau, K., Pai, R., Spiegler, A., Solodkin, A., Jirsa, V., McIntosh, A. R., Ritter, P., & for the Alzheimer’s Disease Neuroimaging Initiative. (2021). Multi-scale brain simulation with integrated positron emission tomography yields hidden local field potential activity that augments machine learning classification of Alzheimer’s disease. https://doi.org/10.1101/2021.02.27.433161

Tzourio-Mazoyer, N., Landeau, B., Papathanassiou, D., Crivello, F., Etard, O., Delcroix, N., Mazoyer, B., & Joliot, M. (2002). Automated anatomical labeling of activations in SPM using a macroscopic anatomical parcellation of the MNI MRI single-subject brain. NeuroImage, 15(1), 273–289. https://doi.org/10.1006/nimg.2001.0978

van Hartevelt, T. J., Cabral, J., Deco, G., Møller, A., Green, A. L., Aziz, T. Z., & Kringelbach, M. L. (2014). Neural plasticity in human brain connectivity: the effects of long term deep brain stimulation of the subthalamic nucleus in Parkinson’s disease. PloS One, 9(1), e86496. https://doi.org/10.1371/journal.pone.0086496

Velliste, M., Kennedy, S. D., Schwartz, A. B., Whitford, A. S., Sohn, J.-W., & McMorland, A. J. C. (2014). Motor cortical correlates of arm resting in the context of a reaching task and implications for prosthetic control. The Journal of Neuroscience: The Official Journal of the Society for Neuroscience, 34(17), 6011–6022. https://doi.org/10.1523/JNEUROSCI.3520-13.2014

Villagrasa, F., Baladron, J., Vitay, J., Schroll, H., Antzoulatos, E. G., Miller, E. K., & Hamker, F. H. (2018). On the Role of Cortex-Basal Ganglia Interactions for Category Learning: A Neurocomputational Approach. The Journal of Neuroscience: The Official Journal of the Society for Neuroscience, 38(44), 9551–9562. https://doi.org/10.1523/JNEUROSCI.0874-18.2018

Visser, S., & Van Gils, S. A. (2014). Lumping Izhikevich neurons. EPJ Nonlinear Biomedical Physics, 2(1), 1–17. https://doi.org/10.1140/epjnbp19

Vitay, J., Dinkelbach, H.Ü., & Hamker, F. H. (2015). ANNarchy: a code generation approach to neural simulations on parallel hardware. Frontiers in Neuroinformatics, 9, 19. https://doi.org/10.3389/fninf.2015.00019

Vitek, J. L., Jain, R., Chen, L., Tröster, A. I., Schrock, L. E., House, P. A., Giroux, M. L., Hebb, A. O., Farris, S. M., Whiting, D. M., Leichliter, T. A., Ostrem, J. L., San Luciano, M., Galifianakis, N., Verhagen Metman, L., Sani, S., Karl, J. A., Siddiqui, M. S., Tatter, S. B.,… Starr, P. A. (2020). Subthalamic nucleus deep brain stimulation with a multiple independent constant current-controlled device in Parkinson’s disease (INTREPID): a multicentre, double-blind, randomised, sham-controlled study. Lancet Neurology, 19(6), 491–501. https://doi.org/10.1016/S1474-4422(20)30108-3

Whitmer, D., de Solages, C., Hill, B., Yu, H., Henderson, J. M., & Bronte-Stewart, H. (2012). High frequency deep brain stimulation attenuates subthalamic and cortical rhythms in Parkinson’s disease. Frontiers in Human Neuroscience, 6, 155. https://doi.org/10.3389/fnhum.2012.00155

Wingeier, B., Tcheng, T., Koop, M. M., Hill, B. C., Heit, G., & Bronte-Stewart, H. M. (2006). Intra-operative STN DBS attenuates the prominent beta rhythm in the STN in Parkinson’s disease. Experimental Neurology, 197(1), 244–251. https://doi.org/10.1016/j.expneurol.2005.09.016

Yu, Y., Wang, X., Wang, Q., & Wang, Q. (2020). A review of computational modeling and deep brain stimulation: applications to Parkinson’s disease. Applied Mathematics and Mechanics. English Edition, 1–22. https://doi.org/10.1007/s10483-020-2689-9

