## Supplementary for "Virtual deep brain stimulation: Multiscale co-simulation of a spiking basal ganglia model and a whole-brain mean-field model with The Virtual Brain"

of the article

*Details of the spiking network model*

For an overview of all variables used in this study, we refer to Supplementary Table 1. For every neuron in a region node modeled as a spiking network, the membrane potential *V* was assumed to follow the equation

$$\begin{aligned}\dot{V} = n_2V^2 + n_1V + n_0 - U/C - g_{AMPA}(V - E_{AMPA}) \\ -g_{GABA}(V_m - E_{GABA}) + I_e + I_{DBS}\end{aligned} \quad (1)$$

$$\dot{U} = a(bV - U), \quad (2)$$

where $a$, $b$, $n_0$, $n_1$ and $n_2$ are region-specific parameters that were taken from literature as in (Maith et al. 2021) (Supplementary Table 2). Specifically, parameters for GPe, GPi, STN were taken from (Thibeault and Srinivasa 2013), for striatum from (Humphries, Wood, and Gurney 2009) and for thalamus taken from the phasic bursting model of (Izhikevich 2004). If the membrane potential exceeds a certain threshold value $V_{th}$, i.e., $V > V_{th}$, a spike is emitted, $V$ is set to a value $c$ and $U$ is incremented by a fixed amount $d$. The conductance follows the equation

$$\dot{g_s} = -g_s/\tau_s + \left[\sum_k \delta(t - t_k)\right]_s, \quad (3)$$

where $s \in \{AMPA, GABA\}$ and $\delta$ is the Kronecker delta. The last term of this previous equation is the increase of the conductance by a fixed amount after each incoming spike $k$ at spike time $t_k$. The equations were taken from (Baladron, Nambu, and Hamker 2019), only $I_{DBS}$ was added.

Similar to previous publications (Maith et al. 2021; Baladron, Nambu, and Hamker 2019), we simulated the above model in ANNarchy (Vitay, Dinkelbach, and Hamker 2015) and applied the Euler method to solve the differential equations (Equations (1)-(3)) with a time step of $0.05ms$.

***Supplementary Table 1: Glossary table of all the used variables of our multiscale model.*** *We list all used variables with a short description, possibly their assigned values and respective unit.*

| variable | assigned value | unit | description |
|---|---|---|---|
| $t$ | - | $ms$ | time |
| $dt$ | $0.1$ | $ms$ | TVB integration time step |
| **Spiking network** | | | |
| $V$ | - | $mV$ | membrane potential |
| $U$ | - | $\frac{mV}{ms}$ | recovery variable |
| $C$ | region-specific, Table 1 | $\frac{1}{pF}$ | membrane capacity |
| $I_e$ | region-specific, Table 1 | $pA$ | region-specific external current |
| $I_{DBS}$ | GPi stimulus: -10<br>monophasic STN: -280<br>biphasic STN: 160 | $pA$ | DBS current |
| $a$ | region-specific, Table 1 | - | rate of recovery of U |
| $b$ | region-specific, Table 1 | - | sensitivity of recovery to subthreshold fluctuations of membrane potential |
| $c$ | region-specific, Table 1 | $mV$ | after-spike reset value of V |
| $d$ | region-specific, Table 1 | $\frac{mV}{ms}$ | after-spike increment of U |
| $n_0$ | region-specific, Table 1 | - | neuron-type specific parameter |
| $n_1$ | region-specific, Table 1 | - | neuron-type specific parameter |
| $n_2$ | region-specific, Table 1 | - | neuron-type specific parameter |
| $V_{th}$ | striatum: 40<br>all others: 30 | $mV$ | spike threshold membrane potential |
| $g_{AMPA}$ | - | $nS$ | AMPA synaptic conductance |
| $g_{GABA}$ | - | $nS$ | GABA synaptic conductance |

| | | | |
|---|---|---|---|
| $E_{AMPA}$ | $0$ | $mV$ | AMPA reversal potential |
| $E_{GABA}$ | $-90$ | $mV$ | GABA reversal potential |
| $\tau_{AMPA}$ | $10$ | $ms$ | AMPA synapse time constant |
| $\tau_{GABA}$ | $10$ | $ms$ | GABA synapse time constant |
| **Mean-field model** | | | |
| $S_{n'}$ | - | - | proportion of open synaptic ion channels |
| $R_{n'}$ | - | $1/s$ | instantaneous spike rate |
| $\tau_{syn}$ | $1000$ | $ms$ | synaptic time scale |
| $\gamma$ | $0.641$ | $s$ | excitatory kinetic parameter |
| $\alpha$ | $2710$ | $nC^{-1}$ | excitatory sigmoidal function parameter |
| $\beta$ | $108$ | $Hz$ | excitatory sigmoidal function parameter |
| $\theta$ | $0.154$ | $s$ | excitatory sigmoidal function parameter |
| $I_{syn_{n'}}$ | - | $nA$ | presynaptic current |
| $I_o$ | $0.3$ | $nA$ | overall effective external input current |
| $w_+$ | $0.9$ | - | local excitatory recurrence |
| $J_N$ | $0.2609$ | $nA$ | excitatory synaptic coupling |
| $a_c$ | $1/256$ | - | linear coupling parameter |
| $b_c$ | $0$ | - | linear coupling parameter |
| $G$ | $15/C_{norm}$ | - | global coupling constant |
| $C_{m'n'}$ | - | - | connection weight from node $n'$ to node $m'$ |
| $\tau_{m'n'}$ | - | $ms$ | delay from node $n'$ to node $m'$ |
| **Interface and connectivity** | | | |
| $w_{n'n}$ | - | - | interface weight / spike weight from proxy node $n'$ to spiking node $n$ |
| $\tau_{n'n}$ | - | $ms$ | spike delay from proxy node $n'$ to spiking node $n$ |
| $t_{jk}$ | - | $ms$ | spike time of neuron $j$ |
| $N_{neurons}$ | - | - | number of neurons in a population |
| $N_{spikes}$ | - | - | number of spikes |

| | | | |
|---|---|---|---|
| $R_{in_n}$ | - | $\frac{1}{ms}$ | input rate |
| $R_{int_n}$ | - | $\frac{1}{ms}$ | integrated input rate |
| $\tau_{int}$ | $100$ | $ms$ | integration time constant |
| $C_{norm}$ | control: $C_{norm} = 0.00068199$ patient: $C_{norm} = 0.000705308$ | - | connectivity weight ratio |
| $w_{norm_{iSN}}$ | control: 0.0382668 patient: 0.0380754 | - | interface factor for iSN |
| $w_{norm_{dSN}}$ | control: 0.0321123 patient: 0.0338178 | - | interface factor for dSN |
| $w_{norm_{STN}}$ | control: 0.311472 patient: 0.2894958 | - | interface factor for STN |
| **DBS stimulus** | | | |
| $a_{DBS}$ | monophasic: -35 biphasic: 20 | $V$ | stimulus amplitude |
| $\kappa$ | $8$ | $pS$ | scaling factor |
| $f$ | monophasic: 120 biphasic: 130 | $Hz$ | stimulus frequency |
| $\delta_{DBS}$ | 0.3 | $ms$ | pulse width (of the first, short and high-amplitude phase for the biphasic stimulus) |

STN: subthalamic nucleus; dSN: striatum, direct striatal spiny projection neurons; iSN: striatum, indirect striatal spiny projection neurons; DBS: deep brain stimulation.

***Supplementary Table 2: Spiking network parameters.*** *Set parameter values for each of the neuronal populations modeled as a spiking network (values taken from (Maith et al. 2021)).*

| Population | a | b | c (mV) | d | C | $I_e$ | $n_0$ | $n_1$ | $n_2$ |
|---|---|---|---|---|---|---|---|---|---|
| Striatum | 0.05 | −20 | −55 | 377 | 50 | 0 | 61.65 | 2.59 | 0.02 |
| GPi | 0.005 | 0.585 | -65 | 4 | 1 | 30 | 140 | 5 | 0.04 |
| GPe | 0.005 | 0.585 | -65 | 4 | 1 | 12 | 140 | 5 | 0.04 |
| STN | 0.005 | 0.265 | -65 | 2 | 1 | 3 | 140 | 5 | 0.04 |
| Thalamus | 0.02 | 0.25 | -65 | 0.05 | 1 | 3.5 | 140 | 5 | 0.04 |
| CxExcit | 0.02 | 0.2 | -72 | 6 | 1 | 50 | 140 | 5 | 0.04 |
| CxInh | 0.02 | 0.2 | -72 | 6 | 1 | 0 | 140 | 5 | 0.04 |

CxExcit: excitatory population of the cortex; CxInh: inhibitory population of the cortex; GPi: internal globus pallidus; GPe: external globus pallidus; STN: subthalamic nucleus.

*Details of the underlying connectivity*

The generation of the atlas from Petersen and colleagues (2019) started from co-registering histological data (Morel 2007; Gallay et al. 2008) and normative MRI-based CIT-168 atlas (Pauli, Nili, and Tyszka 2018) from the Human Connectome Project (Van Essen et al. 2013). As the used histological atlas is focused on the thalamus and BG, additional tracts were approximated by information from non-human studies. The resulting streamlines were manually curated by the neuroanatomists in a holographic augmented-reality interface to correct their three-dimensional approximation interactively and further validated with the underlying histological data. As a result, the connectome by Petersen et al. provides a precise three-dimensional representation of fiber tracts in the human brain and can be used with any existing parcellation to calculate structural connectivity, similar to the calculation from DTI.

*Details about ANNarchy*

Based on the code written by the user, ANNarchy automatically generates optimized C++ code, which can be run on different types of parallel hardware (e.g., on a multi-core system or graphical processing unit). Together with the simulation in automatically generated C++ code and the use of parallel computing, ANNarchy possesses a unique combination of properties, enabling detailed simulations at low computational costs. ANNarchy runs on GNU/Linux and OSX and is open source with freely available documentation and source code at http://annarchy.readthedocs.org and http://bitbucket.org/annarchy/annarchy. An overview of the functions and objects of ANNarchy that were integrated in the TVB-ANNarchy simulation framework is shown in Supplementary Figure 1.

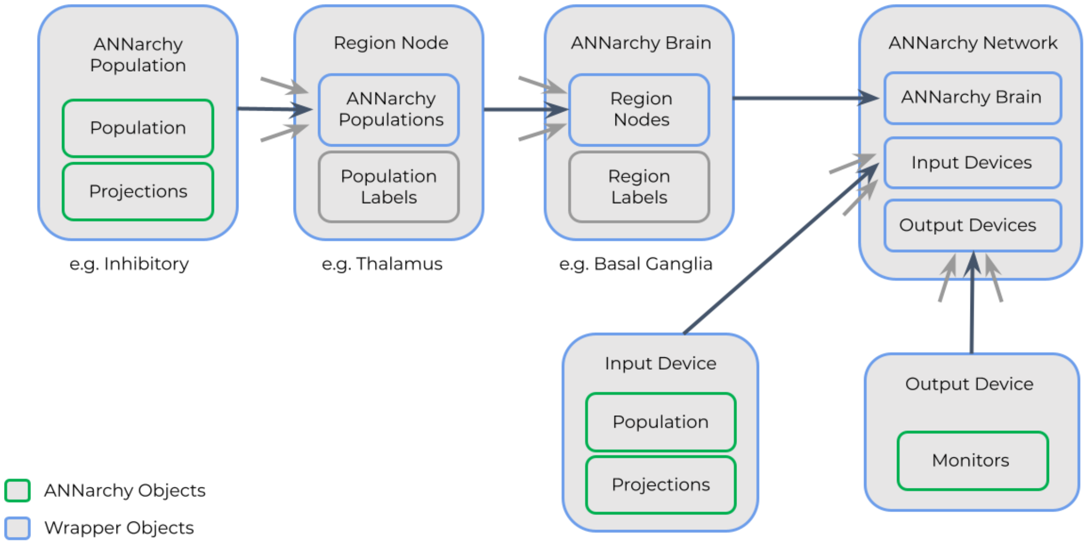


***Supplementary Figure 1: Architecture of the TVB-ANNarchy co-simulation interface.*** *The different existing objects in ANNarchy are represented by gray boxes. The smallest defined objects are called devices and populations. A device ("ANNarchy Device") can be either an input or output device. Input devices contain special ANNarchy populations that can generate stimuli which can be used as inputs to other spiking populations. Output devices wrap around one or several ANNarchy monitors for recording spikes and state variables. A population object called "ANNarchy Population" wraps around a Population or Population View object from ANNarchy and also manages the projections between populations. In our use case, an ANNarchy Population can be an inhibitory population for example. A Region Node is one level above that in the hierarchy. It can contain several ANNarchy Populations and holds their labels. In our case, the thalamus is represented as a Region Node with one population. The ANNarchy Brain is on the next organisational level. It holds all Region Nodes and a mapping to their labels. The ANNarchy Network is the highest-level object. It holds an ANNarchy Brain as well as all Input Device and Output Device objects.Objects that exist in the ANNarchy library are colored in green, in blue are wrappers around them. Gray arrows indicate that several objects of the same type are typically held inside an object higher in the hierarchy, e.g., a Region Node can contain many ANNarchy Populations.*

***Supplementary Table 3: Firing rate validation between the ANNarchy model implementation by (Maith et al., 2021) and our TVB-ANNarchy implementation of the spiking-cortex model.*** *We compared the two spiking network implementations. For this validation, we performed a shorter simulation without noise on the control and patient network data.*

| mean firing rate | ANNarchy model from (Maith et al., 2021) - control | spiking-cortex model inside TVB-ANNarchy - control | ANNarchy model from (Maith et al., 2021) - patient | spiking-cortex model inside TVB-ANNarchy - patient |
|---|---|---|---|---|
| **Cx-E** | 16.0 | 16.0 | 16.0 | 16.0 |
| **Cx-I** | 32.0 | 32.0 | 32.0 | 32.0 |
| **dSN** | 16.4 | 16.6 | 19.2 | 18.9 |
| **iSN** | 15.8 | 15.8 | 15.4 | 15.9 |
| **STN** | 31.4 | 31.3 | 31.9 | 32.0 |
| **GPe** | 34.4 | 34.4 | 35.5 | 35.3 |
| **GPi** | 35.7 | 35.6 | 36.1 | 36.1 |
| **Thal** | 22.7 | 23.1 | 19.9 | 20.3 |

Cx-E: excitatory population of the cortex node; Cx-I: inhibitory population of the cortex node; GPi: internal globus pallidus; GPe: external globus pallidus; STN: subthalamic nucleus; dSN: striatum, direct striatal spiny projection neurons; iSN: striatum, indirect striatal spiny projection neurons; Thal: thalamus.

#### *Mean-field model for the cortical regions*

We used the version of this model that represents each TVB region as one excitatory population. For every region node $n\prime$ (we use the prime notation for nodes modeled only as mean-fields nodes in TVB), the (post-)synaptic gating dynamics $\dot{S_{n\prime}}$ (i.e., $S_{n\prime}$ is the proportion of synapse channels open at any given time) are defined as

$$\dot{S_{n\prime}} = -\frac{1}{\tau_{syn}} S_{n\prime}(t) + (1 - S_{n\prime}(t))\gamma R_{n\prime}(t),$$

where $\tau_{syn} = 1000ms$ is the time scale of the synapse and $R_{n\prime}(t)$ is the postsynaptic firing rate given by

$$R_{n\prime}(t) = \frac{\alpha I_{syn_{n\prime}}(t) - \beta}{1 - e^{-\theta(\alpha I_{syn_{n\prime}}(t) - \beta)}},$$

which is a sigmoidal activation function of the presynaptic input current $I_{syn_{n\prime}}$. The total presynaptic current is given by

$$I_{syn_{n\prime}}(t) = I_o + w_+ J_N S_{n\prime}(t) + a_c G J_N \sum_{m\prime \neq n\prime} C_{m\prime n\prime} S_{m\prime}(t - \tau_{m\prime n\prime}).$$

The variables $C_{m\prime n\prime}$ and $\tau_{m\prime n\prime}$ define the weight and delay for the connection from region node $n\prime$ to region node $m\prime$, respectively, whereby the sum runs over all pairwise combinations. The connectivity weights were additionally scaled by the global coupling constant $G$ and the parameter $a_c$ of TVB's linear coupling function (of the form $a_c x + b_c$, in which we set the default parameters $b_c = 0$ and $a_c = 1/256$). The parameters were determined as in (Deco et al. 2013) (Supplementary Table 1).

*Details of the spike train generation for the coupling from TVB to ANNarchy*

For the correlated spike trains' generation, TVB "proxy" nodes are modeled in ANNarchy, as populations Poisson-like spiking neurons whose population rate *x* varies following a stochastic differential equation

$$\frac{dx}{dt} = \frac{\mu - x}{\tau} + \sigma \frac{\xi}{\sqrt{\tau}},$$

where $\xi$ is a random variable. Thus, the population rate $x$ randomly varies around $\mu$ over time, with an amplitude determined by $\sigma$ and a speed determined by $\tau$. To avoid that $x$ becomes negative, the values of $\mu$ and $\sigma$ are computed from a rectified Gaussian distribution, parameterized by the desired population rate $R_n$, the desired correlation strength *corr = 0.3*, and the time constant $\tau = 10ms$. In our case, the rate was determined by TVB input at each TVB time step and $\mu$ and $\sigma$ were automatically recomputed by the ANNarchy class *HomogeneousCorrelatedSpikeTrains* (Brette 2009). The correlation parameter of the *HomogeneousCorrelatedSpikeTrains* ANNarchy spike generator was set to $corr = 0.3$ in all cases and models, after visual inspection and comparison of the generated spike trains to the spike trains of the noisy Izhikevich excitatory spiking cortex population used in Maith et al. (2021), which was the driver of the spiking network in that study (Maith et al. 2021).

For the "trial and error" simulations for determining the value of $G$, we started with setting initial conditions of all $S_{n\prime} = 0.0$ and $R_{n\prime} = 0.0$. For every subsequent simulation, we were using the mean state variables' values of the last 100 ms of the previous simulation as initial conditions. The mean firing rate that we were trying to approximate was also computed for those last 100 ms. Once the equilibrium was approximated and the mean firing rate was in the interval *[15, 18] Hz*, we set the $G$ value accordingly and stored the initial conditions. For the *10* repetitive co-simulations for the results of Supplementary Table 4, we selected initial conditions randomly in the neighborhood of the above original vector of initial conditions as it is explained in the main text.

Small differences in the rates obtained for multiscale model's co-simulations are to be expected, especially for the firing rate of the thalamus, which depends a lot on the spikes' correlations among neurons of the populations that couple to it directly and indirectly. In that respect, please note that (a) the weighted superposition of the activity of many TVB "proxy" nodes results in an effective driving dynamics of a quite different autocorrelation profile than that of a single noisy spiking-cortex node, and (b) we have set the same, undifferentiated, value for the correlation of the driving spike generators for both subjects, as explained above.

*Details of the applied STN stimuli*

The monophasic stimulus

$$I_{DBS} = a_{DBS} \cdot \kappa \cdot H\left(sin\left(2\pi ft\right)\right) \cdot \left(1 - H\left(sin\left(2\pi f(t + \delta_{DBS})\right)\right)\right)$$

is adapted from (Michmizos and Nikita 2011), where $H$ is the Heaviside function, $f$ is the frequency in $Hz$, $a_{DBS}$ is the amplitude of the stimulus in *V*, $\kappa = 8$ is a scaling factor, $\delta_{DBS}$ is the pulse width and $t$ is the time in seconds. The biphasic stimulus was defined as

$$I_{DBS} = 1.1 \cdot a_{DBS} \cdot \kappa \cdot H\left(sin\left(2\pi\frac{f}{1000}t\right)\right) \cdot \left(1 - H\left(sin\left(2\pi\frac{f}{1000}(t - \delta_{DBS})\right)\right)\right)$$
$$-0.1 \cdot a_{DBS} \cdot \kappa \cdot H\left(sin\left(2\pi\frac{f}{1000}t\right)\right) \cdot \left(1 - H\left(sin\left(2\pi\frac{f}{1000}(t - (11 \cdot \delta_{DBS}))\right)\right)\right)$$

similar to (Liu et al. 2020). Here, the pulse width $\delta_{DBS}$ is the pulse width of the first, short and high-amplitude phase. The second phase of the biphasic stimulus is designed to be 10 times as long and has *1/10^th^* of the amplitude (Figure 6A). The parameters of the monophasic and the biphasic stimuli are listed in Supplementary Table 1.

***Supplementary Table 4: Regions of the left hemisphere included in the connectome used for simulations.*** *The regions 1-5 are the regions of the basal ganglia network. The other regions are regions from the AAL atlas.*

| Region number | Abbreviation of the region | Full region name |
|---|---|---|
| 1 | GPe | Globus pallidus externus |
| 2 | GPi | Globus pallidus internus |
| 3 | STN | Subthalamic nucleus |
| 4 | Striatum | Striatum |
| 5 | Thal | Thalamus |
| 6 | Precentral | Precentral gyrus |
| 7 | Frontal_Sup_2 | Superior frontal gyrus |
| 8 | Frontal_Mid_2 | Middle frontal gyrus |
| 9 | Frontal_Inf_Oper | Inferior frontal gyrus, opercular part |

| | | |
|---|---|---|
| 10 | Frontal_Inf_Tri | Inferior frontal gyrus, triangular part |
| 11 | Frontal_Inf_Orb_2 | Inferior frontal gyrus, orbital part |
| 12 | Rolandic_Oper | Rolandic operculum |
| 13 | Supp_Motor_Area | Supplementary motor area |
| 14 | Olfactory | Olfactory cortex |
| 15 | Frontal_Sup_Medial | Superior frontal gyrus, medial |
| 16 | Frontal_Med_Orb | Superior frontal gyrus, medial |
| 17 | Rectus | Gyrus rectus |
| 18 | OFCmed | Medial orbital gyrus |
| 19 | OFCant | Anterior orbital gyrus |
| 20 | OFCpost | Posterior orbital gyrus |
| 21 | OFClat | Lateral orbital gyrus |
| 22 | Insula | Insula |
| 23 | Cingulate_Ant | Anterior cingulate & paracingulate gyri |
| 24 | Cingulate_Mid | Middle cingulate & paracingulate gyri |
| 25 | Cingulate_Post | Posterior cingulate gyrus |
| 26 | Hippocampus | Hippocampus |
| 27 | ParaHippocampal | Parahippocampal gyrus |
| 28 | Amygdala | Amygdala |
| 29 | Calcarine | Calcarine fissure and surrounding cortex |
| 30 | Cuneus | Cuneus |

| | | |
|---|---|---|
| 31 | Lingual | Lingual gyrus |
| 32 | Occipital_Sup | Superior occipital gyrus |
| 33 | Occipital_Mid | Middle occipital gyrus |
| 34 | Occipital_Inf | Inferior occipital gyrus |
| 35 | Fusiform | Fusiform gyrus |
| 36 | Postcentral | Postcentral gyrus |
| 37 | Parietal_Sup | Superior parietal gyrus |
| 38 | Parietal_Inf | Inferior parietal gyrus, excluding supramarginal and angular gyri |
| 39 | SupraMarginal | Supramarginal gyrus |
| 40 | Angular | Angular gyrus |
| 41 | Precuneus | Precuneus |
| 42 | Paracentral_Lobule | Paracentral lobule |
| 43 | Heschl | Heschl gyrus |
| 44 | Temporal_Sup | Superior temporal gyrus |
| 45 | Temporal_Pole_Sup | Temporal pole: superior temporal gyrus |
| 46 | Temporal_Mid | Middle temporal gyrus |
| 47 | Temporal_Pole_Mid | Temporal pole: middle temporal gyrus |
| 48 | Temporal_Inf | Inferior temporal gyrus |
| 49 | Cerebelum_Crus1 | Crus I of cerebellar hemisphere |
| 50 | Cerebelum_Crus2 | Crus II of cerebellar hemisphere |

| | | |
|---|---|---|
| 51 | Cerebelum_3 | Lobule III of cerebellar hemisphere |
| 52 | Cerebelum_4_5 | Lobule IV, V of cerebellar hemisphere |
| 53 | Cerebelum_6 | Lobule VI of cerebellar hemisphere |
| 54 | Cerebelum_7b | Lobule VIIB of cerebellar hemisphere |
| 55 | Cerebelum_8 | Lobule VIII of cerebellar hemisphere |
| 56 | Cerebelum_9 | Lobule IX of cerebellar hemisphere |
| 57 | Cerebelum_10 | Lobule X of cerebellar hemisphere |

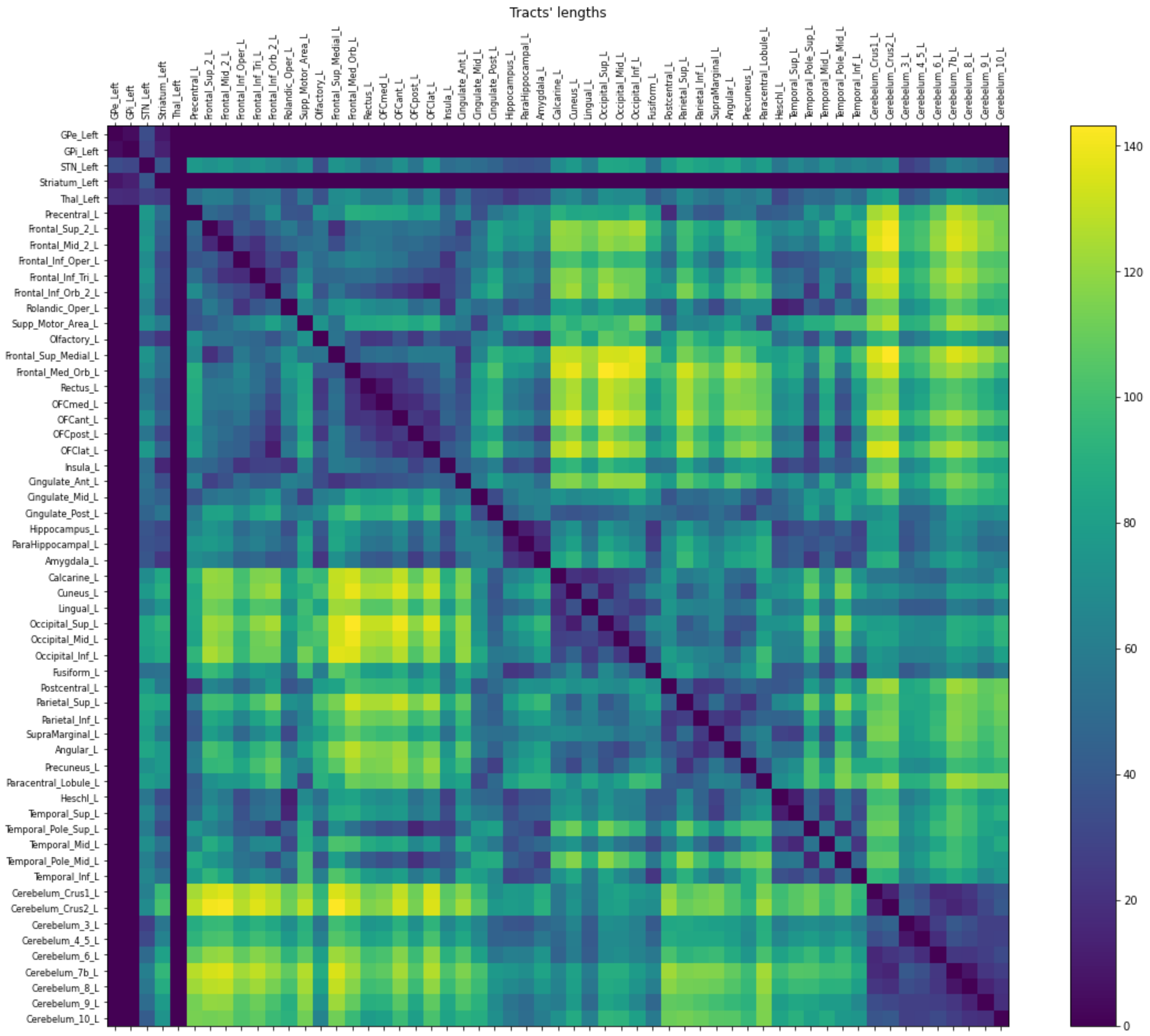


***Supplementary Figure 2: Tract length matrix used for simulations.*** *We approximated the tract lengths among regions by the Euclidean distance between the three-dimensional anatomical center coordinates of each region.*

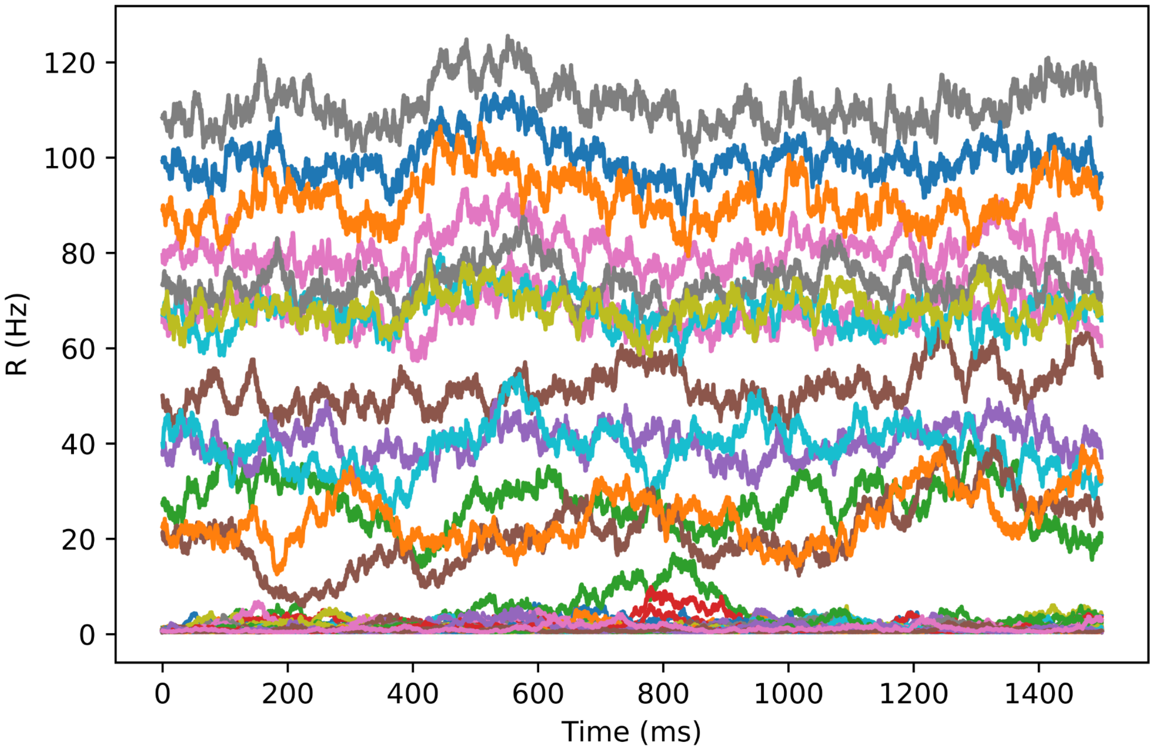


***Supplementary Figure 3: Characteristic time series of the rate state variable $R$ of all TVB nodes from a co-simulation.*** *Each colored time series represents the time series of the state variable R, the firing rate, of one of the TVB nodes. The dynamics consists of random fluctuations (due to additive white noise of standard deviation $10^{-4}$) around an equilibrium point, which is determined by the couplings among the nodes on the basis of the structural connectome (weights and delays) and the global coupling scaling $G$.*

***Supplementary Table 5: Table of obtained firing rates for the spiking-cortex model for patient and control network, in both resting-state and stimulus condition.*** *Mean firing rates averaged over the last 1000ms of each simulation and averaged over the 10 repetition simulations in case of the TVB-cortex model (standard deviation of these mean firing rates over the 10 repetition simulations in case of the TVB-cortex model).*

| **mean firing rate** | **resting-state control** | | **resting-state patient** | | **GPi-DBS patient** | | **STN-DBS biphasic patient** | | **STN-DBS monophasic patient** | |
|---|---|---|---|---|---|---|---|---|---|---|
| | **spiking -cortex model** | **TVB-cortex model** | **spiking -cortex model** | **TVB-cortex model** | **spiking -cortex model** | **TVB-cortex model** | **spiking -cortex model** | **TVB-cortex model** | **spiking -cortex model** | **TVB-cortex model** |
| **Cx-E** | 14.1 | - | 14.1 | - | 14.1 | - | 14.1 | - | 14.1 | - |
| **dSN** | 16.2 | 16.2 (0.1) | 19.9 | 20.7 (0.2) | 22.8 | 23.4 (0.3) | 22.8 | 22.3 (0.4) | 21.5 | 22.7 (0.2) |
| **iSN** | 10.8 | 10.3 (0.1) | 9.5 | 8.9 (0.1) | 11.4 | 11.0 (0.2) | 10.9 | 10.3 (0.3) | 10.5 | 10.6 (0.1) |
| **STN** | 26.9 | 27.4 (0.1) | 29.4 | 30.2 (0.1) | 29.5 | 30.4 (0.1) | 42.9 | 45.9 (0.2) | 28.0 | 27.9 (0.2) |
| **GPe** | 35.3 | 34.8 (0.3) | 35.7 | 35.5 (0.1) | 35.6 | 35.4 (0.2) | 40.3 | 40.7 (0.5) | 36.3 | 35.8 (0.2) |
| **GPi** | 34.2 | 34.5 (0.1) | 34.6 | 34.6 (0.1) | 30.3 | 30.5 (0.1) | 38.9 | 39.6 (0.3) | 35.2 | 35.9 (0.1) |
| **Thal** | 17.4 | 19.4 (0.2) | 12.8 | 12.6 (0.3) | 20.4 | 19.3 (0.5) | 17.9 | 16.5 (0.9) | 16.9 | 17.2 (0.3) |

Cx-E: excitatory population of the cortex node; GPi: internal globus pallidus; GPe: external globus pallidus; STN: subthalamic nucleus; dSN: striatum, direct striatal spiny projection neurons; iSN: striatum, indirect striatal spiny projection neurons; Thal: thalamus.

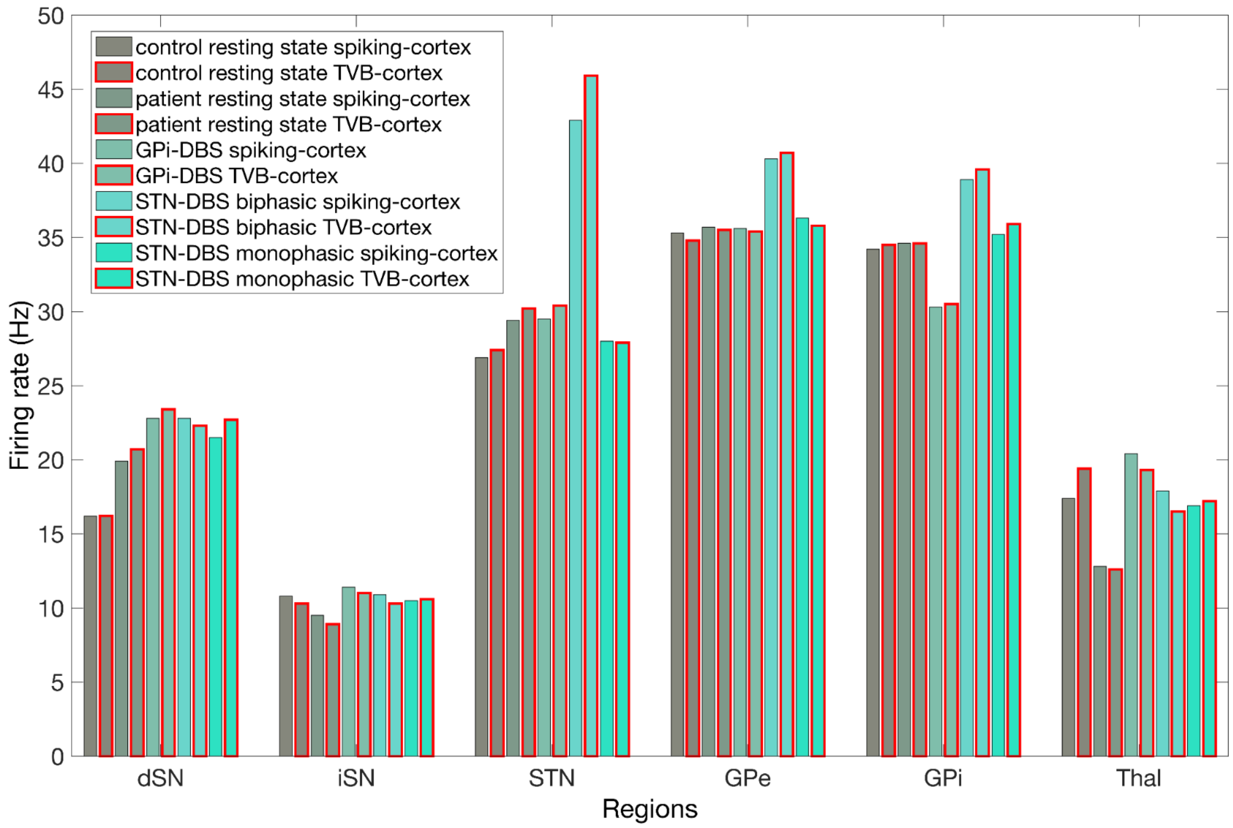


***Supplementary Figure 4: Average firing rates obtained by different simulations of the TVB-cortex and the spiking-cortex model.*** *For each of the six spiking regions, each bar represents either the resting-state condition for the control, the patient or one of the three virtual DBS simulations, i.e., GPi-DBS, STN-DBS applying a biphasic and a monophasic stimulus. For each region, the first, third, fifth, seventh and ninth bars represent the average firing rate of the spiking-cortex simulation, the second, fourth, sixth, eighth and tenth bars (with red outline) represent the average firing rate of the TVB-cortex simulations. The heights of the bars represent the firing rate (in Hz) averaged over the last 1000 ms of the respective simulation and over the 10 simulation repetitions for the TVB-cortex simulations. GPi: internal globus pallidus; GPe: external globus pallidus; STN: subthalamic nucleus; dSN: striatum, direct striatal spiny projection neurons; iSN: striatum, indirect striatal spiny projection neurons; Thal: thalamus.*

### References

Baladron, Javier, Atsushi Nambu, and Fred H. Hamker. 2019. "The Subthalamic Nucleus-External Globus Pallidus Loop Biases Exploratory Decisions towards Known Alternatives: A Neuro-Computational Study." *The European Journal of Neuroscience* 49 (6): 754–67.

Brette, Romain. 2009. "Generation of Correlated Spike Trains." *Neural Computation* 21 (1): 188–215.

Deco, Gustavo, Adrián Ponce-Alvarez, Dante Mantini, Gian Luca Romani, Patric Hagmann, and Maurizio Corbetta. 2013. "Resting-State Functional Connectivity Emerges from Structurally and Dynamically Shaped Slow Linear Fluctuations." *The Journal of Neuroscience: The Official Journal of the Society for Neuroscience* 33 (27): 11239–52.

Gallay, Marc N., Daniel Jeanmonod, Jian Liu, and Anne Morel. 2008. "Human Pallidothalamic and Cerebellothalamic Tracts: Anatomical Basis for Functional Stereotactic Neurosurgery." *Brain Structure & Function* 212 (6): 443–63.

Humphries, Mark D., Ric Wood, and Kevin Gurney. 2009. "Dopamine-Modulated Dynamic Cell Assemblies Generated by the GABAergic Striatal Microcircuit." *Neural Networks: The Official Journal of the International Neural Network Society* 22 (8): 1174–88.

Izhikevich, Eugene M. 2004. "Which Model to Use for Cortical Spiking Neurons?" *IEEE Transactions on Neural Networks / a Publication of the IEEE Neural Networks Council* 15 (5): 1063–70.

Liu, Chen, Ge Zhao, Jiang Wang, Hao Wu, Huiyan Li, Chris Fietkiewicz, and Kenneth A. Loparo. 2020. “Neural Network-Based Closed-Loop Deep Brain Stimulation for Modulation of Pathological Oscillation in Parkinson’s Disease.” *IEEE Access* 8: 161067–79.

Maith, Oliver, Francesc Villagrasa Escudero, Helge Ülo Dinkelbach, Javier Baladron, Andreas Horn, Friederike Irmen, Andrea A. Kühn, and Fred H. Hamker. 2021. “A Computational Model-Based Analysis of Basal Ganglia Pathway Changes in Parkinson’s Disease Inferred from Resting-State fMRI.” *The European Journal of Neuroscience* 53 (7): 2278–95.

Michmizos, Kostis P., and Konstantina S. Nikita. 2011. “Addition of Deep Brain Stimulation Signal to a Local Field Potential Driven Izhikevich Model Masks the Pathological Firing Pattern of an STN Neuron.” *Conference Proceedings: ... Annual International Conference of the IEEE Engineering in Medicine and Biology Society. IEEE Engineering in Medicine and Biology Society. Conference* 2011: 7290–93.

Morel, Anne. 2007. *Stereotactic Atlas of the Human Thalamus and Basal Ganglia*. CRC Press.

Pauli, Wolfgang M., Amanda N. Nili, and J. Michael Tyszka. 2018. “A High-Resolution Probabilistic in Vivo Atlas of Human Subcortical Brain Nuclei.” *Scientific Data* 5 (April): 180063.

Thibeault, Corey M., and Narayan Srinivasa. 2013. “Using a Hybrid Neuron in Physiologically Inspired Models of the Basal Ganglia.” *Frontiers in Computational Neuroscience* 7 (July): 88.

Van Essen, David C., Stephen M. Smith, Deanna M. Barch, Timothy E. J. Behrens, Essa Yacoub, Kamil Ugurbil, and WU-Minn HCP Consortium. 2013. “The WU-Minn Human Connectome Project: An Overview.” *NeuroImage* 80 (October): 62–79.

Vitay, Julien, Helge Ü. Dinkelbach, and Fred H. Hamker. 2015. “ANNarchy: A Code Generation Approach to Neural Simulations on Parallel Hardware.” *Frontiers in Neuroinformatics* 9 (July): 19.
